# Left-right cortical interactions drive intracellular pattern formation in the ciliate *Tetrahymena*

**DOI:** 10.1101/2024.12.18.629246

**Authors:** Chinkyu Lee, Ewa Joachimiak, Wolfgang Maier, Yu-Yang Jiang, Karl F. Lechtreck, Eric S. Cole, Jacek Gaertig

**Affiliations:** Department of Cellular Biology, University of Georgia, Athens, GA 30602, USA; Nencki Institute of Experimental Biology of Polish Academy of Sciences, 02-093 Warsaw, Poland; Bioinformatics, University of Freiburg, 79110 Freiburg, Germany; Biology Department, St. Olaf College, Northfield, MN 55057, USA

## Abstract

In ciliates, cortical organelles are positioned at precise locations along two polarity axes: anterior-posterior and circumferential (lateral). We explored the poorly understood mechanism of circumferential patterning, which generates left-right asymmetry. The model ciliate *Tetrahymena* has a single anteriorly-located oral apparatus. During cell division, a single new oral apparatus forms near the equator of the parental cell and along the longitude of the parental organelle. Cells homozygous for *hypoangular 1* (*hpo1*) alleles, assemble multiple oral apparatuses positioned either to the left or right flanking the normal oral longitude. We identified *HPO1* as a gene encoding an ARMC9-like protein. Hpo1 colocalizes with ciliary basal bodies, forming a bilateral concentration gradient with the high point on the cell’s right side and a sharp drop-off that marks the longitude at which oral development initiates on the ventral side. Hpo1 acts to exclude oral development from the cell’s right side. Hpo1 interacts with the Beige-Beach domain protein Bcd1, a cell’s left side-enriched factor, whose loss also confers formation of multiple oral apparatuses. A loss of both Hpo1 and Bcd1 is lethal and profoundly disrupts both positioning and organization of the forming oral apparatus (including its internal left-right polarity). We conclude that in ciliates, the circumferential/chiral patterning involves gradient-forming factors that are concentrated on either the cell’s right or left side and that the two sides of the cortex interact to create boundary effects that induce, position and shape developing cortical organelles.

## Introduction

Eukaryotic cells typically have a prominent anterior-posterior (also called front-to-rear or apical-basal) polarity. In addition, some cell types, including cells that move while adhering to a substrate or single-cell metazoan embryos, also have a distinguishable dorsal-ventral polarity axis. Moreover, some cells display chiral asymmetries in reference to the anterior-posterior axis. Ciliated protists have features that are favorable for exploring how multiple polarities and chiral asymmetries are generated in the cell. In the genetic model ciliate *Tetrahymena thermophila* cortical organelles occupy specific positions along the anterior-posterior axis and around the cell circumference (Fig. 1A-1). All cortical structures are duplicated during a type of binary fission called “tandem duplication”, in the course of which a single parental cell is subdivided into two daughter cells arranged head-to-tail (Fig. 1A). The anterior daughter cell assembles new posterior organelles while the posterior daughter cell assembles new anterior organelles. A new oral apparatus (oral primordium) forms on the ventral surface near the future anterior end of the posterior daughter cell (Fig. 1A-2). Remarkably, the primordium almost always assembles in association with basal bodies of a single specific ciliary row (row 0, Fig. 1B). The anterior-posterior position of the oral primordium is regulated by a set of conserved cortical proteins, (including components of the Hippo signaling pathway) that appear to act by marking cortical domains where organelle assembly is disallowed (reviewed by ^1^). The principles of lateral positioning (around the circumferential axis) are less understood but could be exceptionally revealing as the responsible mechanism must generate precise coordinates in the absence of structural landmarks available for the anterior-posterior positioning (such as cell ends). Furthermore, studies in ciliates may provide insights into mechanisms that generate chiral asymmetries. In multicellular models, mutations in cytoskeletal proteins (including myosin, α-tubulin and β-tubulin) can invert the left-right positioning of body organs (^2-5^, reviewed in ^6^). Intriguingly in ciliates, alterations of left-right organelle positioning can be caused by mutations in non-cytoskeletal proteins (including a protein associated with vesicle trafficking and a kinase ^7,8^) or by trauma that alters the cortical organization in a way that is epigenetically heritable across multiple cell generations ^9,10^.

**Figure 1.**
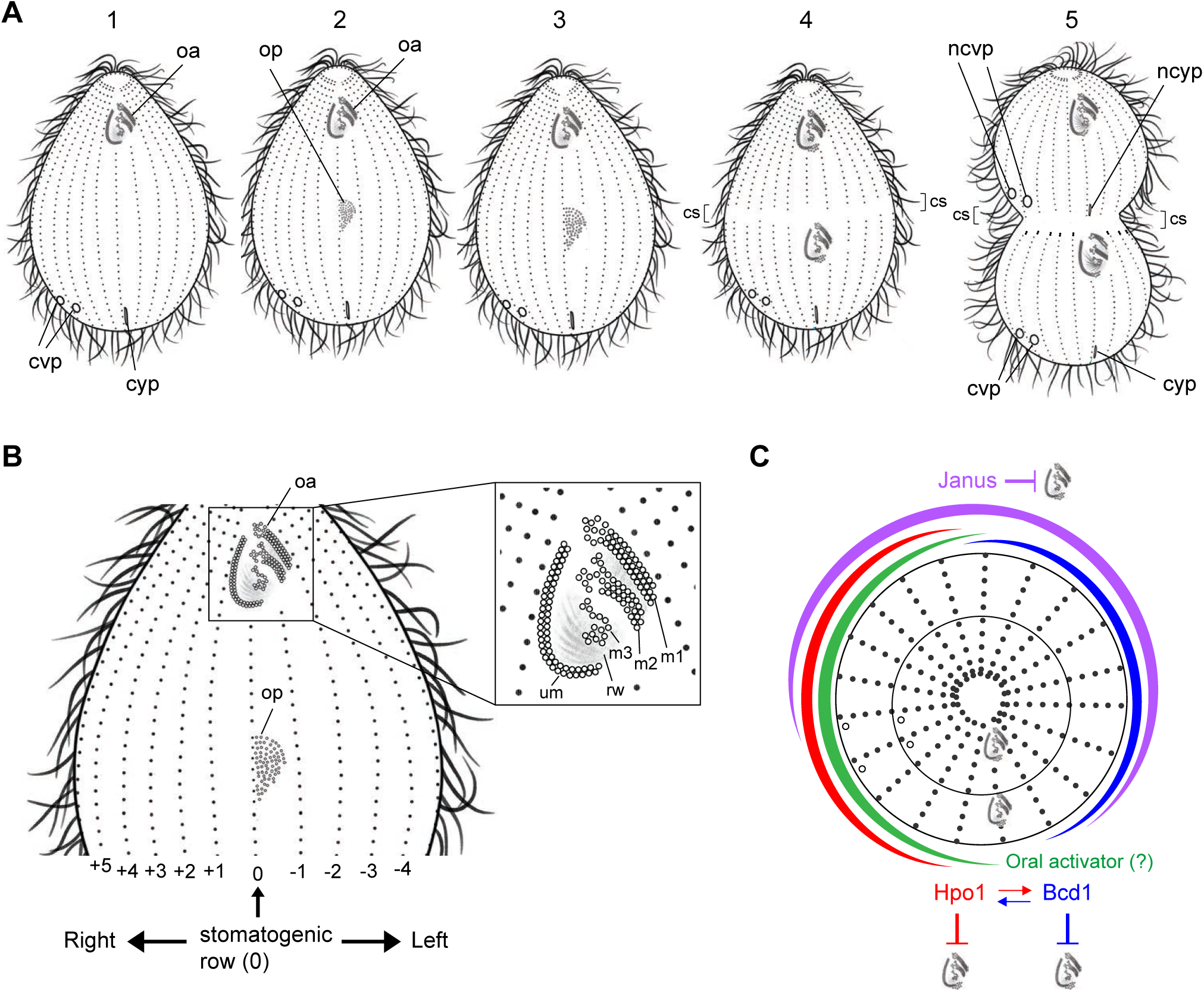
Circumferential positioning In *Tetrahymena*. (A) Cell cycle stages of *T. thermophila* with emphasis on cortical development. Note that during cell division, new organelles form at same longitudes (same ciliary rows) as old organelles. (B) A detailed view of a stage 2 cell presents the row numbering method. (C) A multi-domain model for circumferential positioning in a ciliate. Abbreviations: oa, oral apparatus; op, oral primordium; cvp, contractile vacuole pore; ncvp, new contractile vacuole pore; m1,m2,m3 oral M (membranelle) rows; um, oral undulating membrane row.

*Tetrahymena thermophila* is a fast growing ciliate that is widely used as a genetic model with both reverse ^11^ and forward ^12^ genetics strategies. A library of unique *Tetrahymena* mutants with defects in cortical patterning has been developed and extensively characterized by Joseph Frankel and colleagues (University of Iowa) ^13,14^. The mutants of interest here are those in which the principal defect is in the positioning of organelles around the circumferential axis. The *hpo1* and *bcd1* alleles ^7,15,16^ exhibit multiple oral primordia assembled at incorrect circumferential locations. Recently, Cole and colleagues identified the *BCD1* gene product as a Beige-BEACH domain protein orthologous to the mammalian NBEA and Rugose of *D. melanogaster*. Bcd1 is enriched on the cell’s left side and appears to act by regulating the balance between delivery and retrieval of organelle assembly components through endocytosis and exocytosis, to control the size of cortical domains competent for organelle formation ^7^. Here, we used comparative next generation sequencing (NGS) to identify *HPO1* as a gene encoding a protein similar to ARMC9, a highly conserved ciliary distal tip protein. Remarkably, Hpo1 is enriched on the *Tetrahymena* cell’s right side where it forms a bilateral concentration gradient that diminishes both dorsally and ventrally. Oral development occurs at positions at which Hpo1 levels drop abruptly. Our data suggest that Hpo1 functions as a bidirectional repressor excluding oral morphogenesis from the cell’s right lateral side. We also reveal that together, the right-side-biased Hpo1 and the left-side-biased Bcd1 are required for cell viability and that mutants lacking both proteins display diverse and severe patterning defects including local defects in the organization of cortical structures. We propose that in a ciliate, circumferential patterning involves multiple cortical subdomains formed by bilateral gradients of pattern regulators that interact with each other and act by generating cortical boundaries that define sites of organelle assembly.

## Results

### Hpo1 is an ARMC9-like protein, TTHERM_001276421

*Tetrahymena* cells have two permanent polarity axes: anterior-posterior (AP) and circumferential (C). AP polarity is reflected by an asymmetric placement of major cortical organelles: the oral apparatus (OA) near the anterior cell end, the contractile vacuole pores (CVPs) and the cytoproct near the posterior cell end (Fig. 1A-1). C axis polarity is revealed by asymmetric lateral positions of cortical organelles. While the cytoproct is located at the same longitude as the OA, the CVPs are located on the cell’s right lateral side (Fig. 1A-1). During cell division new organelles form at the same cell longitudes as the preexisting organelles. The new OA (oral primordium or OP) assembles initially as a group of ciliary basal bodies (BBs) forming at a subequatorial AP position in the middle of the cell’s ventral side, precisely on the left side of row 0 (Fig. 1A-2). BBs proliferate and form four oral ciliary rows (UM on the right and M1, M2 and M3 on the left) (Fig. 1A-4, 1B). In an advanced stage of oral development, a fission zone bisects all somatic ciliary rows anteriorly to the OP (Fig. 1A-3-5). The new CVPs and the new cytoproct form at the posterior ends of the anterior half-rows (Fig. 1A-5). The cell completes nuclear divisions and undergoes cytokinesis (Fig. 1A-5).

We adopted the longitudinal row numbering scheme after Cole and colleagues ^15^. The right postoral (stomatogenic) row is designated as row 0 (Fig. 1B). The remaining rows are numbered as +1 or higher when moving to the cell’s right side or -1 or lower when moving to the cell’s left side (Fig. 1B). *hpo1* (*hypoangular 1*) alleles disturb the positioning of oral development along the C axis ^16^. While in the wild type a single OP forms near the left side of row 0, in the *hpo1* mutants, multiple OPs form near several adjacent ventral rows located to either the left (rows -1, -2…) or right (rows +1, +2…) of the stomatogenic row 0 (Fig. 2B,E compare to Fig. 2A,D; Fig. 2H). One complication is that in some *hpo1* mutant cells the oral apparatus is wider than normal, which is correlated with the presence of 3 post-oral rows. In such cases we designated the middle postoral row as 0 ^16^. As described by Frankel and colleagues ^16^, in the *hpo1-3* mutants, the extra OPs form more frequently to the right of row 0 (Fig. 2B,E,H). However, when multiple OPs were present, they were located near adjacent rows, typically including row 0 or a row in the proximity of row 0. Thus, in the *hpo1* mutants, the OP positions are not random. The *hpo1* phenotype can be described as an inability to focus to OP position to a single ventral row. Also as described ^16^, the severity of the *hpo1* allele phenotypes increases at 38°C as compared to the standard temperature 30°C (Fig. 2H). As oral development progresses, adjacent OPs frequently fuse into one oral field and consequently most mutant cells have a single, mature OA (OA in Fig. 2B,E). Intriguingly, the *hpo1* alleles also reduce the number of CVPs per cell: while most wild-type cells have two CVPs (near two adjacent rows on the right side, usually +4 and +5), *hpo1* mutant cells often have only one CVP ^16^, (Fig. S1B compare to Fig. S1A; Fig. S1H).

**Figure 2.**
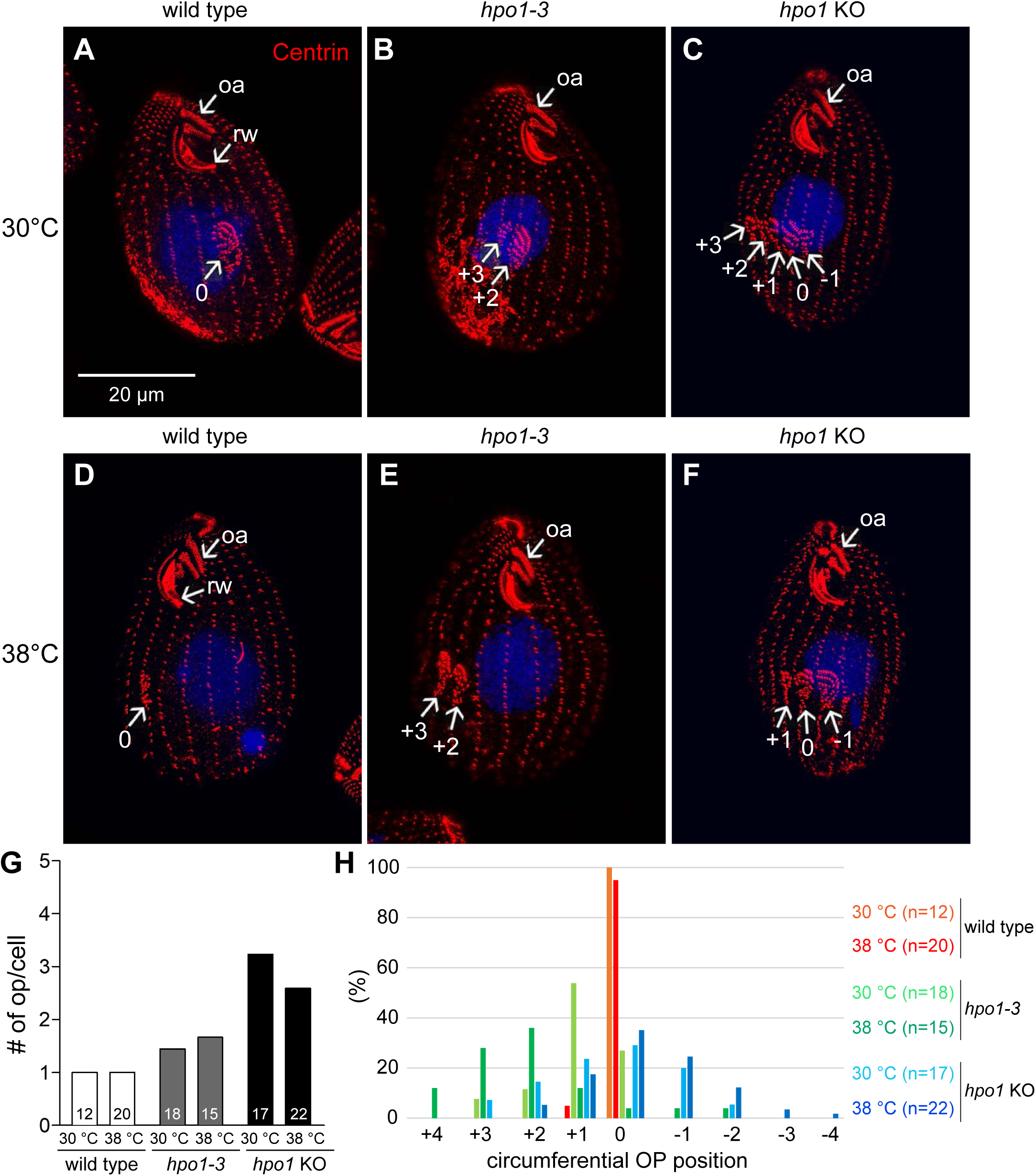
*hpo1* alleles disturb the patterning on the circumferential axis by conferring an excessive number of oral primordia and their lateral displacement. (A-F) SR-SIM images of cells labeled with the 20H5 anti-centrin antibody (red) and DAPI (blue). The cells were cultured overnight at either 30°C (A-C) or 38°C (D-F). (G,H) The graphs document an increase in the number of oral primordia (G) and their lateral displacement (H) conferred by the *hpo1* alleles. Abbreviations: oa, oral apparatus; rw, ribbed wall of microtubules. The numbers mark the C positions of ventral rows with OPs.

We used comparative NGS to map genomic variants that co-segregate with each of the four available *hpo1* alleles (*hpo1-1*, *hpo1-2*, *hpo1-3* and *hpo1-4*). Linkage peaks were consistently detected on the micronuclear chr3 around ∼5 Mb (see the data obtained for *hpo1-3* in Fig. 3A), in agreement with the previous assignment of *hpo1-1* to chr3 using complementation tests in crosses between mutant homozygotes and nullisomic strains lacking specific micronuclear chromosomes ^16^. Within the linkage peak region, we located two homozygous variants: chr3:5363773 G/A in *hpo1-1* and *hpo1-4* and chr3:5364108 T/C in *hpo1-2* and *hpo1-3* genomes, respectively. Both variants are located within the same protein-coding gene, *TTHERM_001276421*. BlastP searches identified TTHERM_001276421 homologs in diverse ciliate species including another oligohymenophoran *Paramecium tetraurelia* (GSPATG00001303001 and 6 paralogs), and more evolutionarily-distant ciliates: the hypotrich *Oxytricha fallax* (g7224, g191167 and g3871) and the heterotrich *Stentor coeruleus* (SteCoe_23425, SteCoe_24205). While BlastP searches failed to identify proteins with significant homology outside of the ciliate phylum, the domain organization of TTHERM_001276421 resembles that of the conserved ciliary tip protein ARMC9, whose mutations cause Joubert syndrome ^17-19^. Both ARMC9 and TTHERM_001276421 have the same set of protein domains arranged in the same order (Fig. 3B,B’). The N-terminal one-third of TTHERM_001276421 is classified by Interpro as the ARMC9 domain (IPRO040369), within which there is a LisH domain (IPR006594), and two coiled-coil regions (Fig. 3B,B’). The C-terminal region contains an ARM-like domain (IPRO11989) (Fig. 3B,B’). Thus, it appears that TTHERM_001276421 is a ciliate lineage-specific ARMC9-like protein. The chr3:5363773 G/A variant (*hpo1-1*, *hpo1-4*) results in the S236N amino acid substitution, while the chr3:5364108 T/C variant (*hpo1-2*, *hpo1-3*) results in the F318S substitution, both within the ARM-like domain. In agreement with our findings, homozygotes for *hpo1-2* and *hpo1-3* were reported to have a similar temperature-sensitive phenotype, less severe than that of the *hpo1-1* and *hpo1-4* homozygotes (^16^, and information deposited at the *Tetrahymena* Stock Center (https://tetrahymena.vet.cornell.edu/display.php?stockid=SD01466).

**Figure 3.**
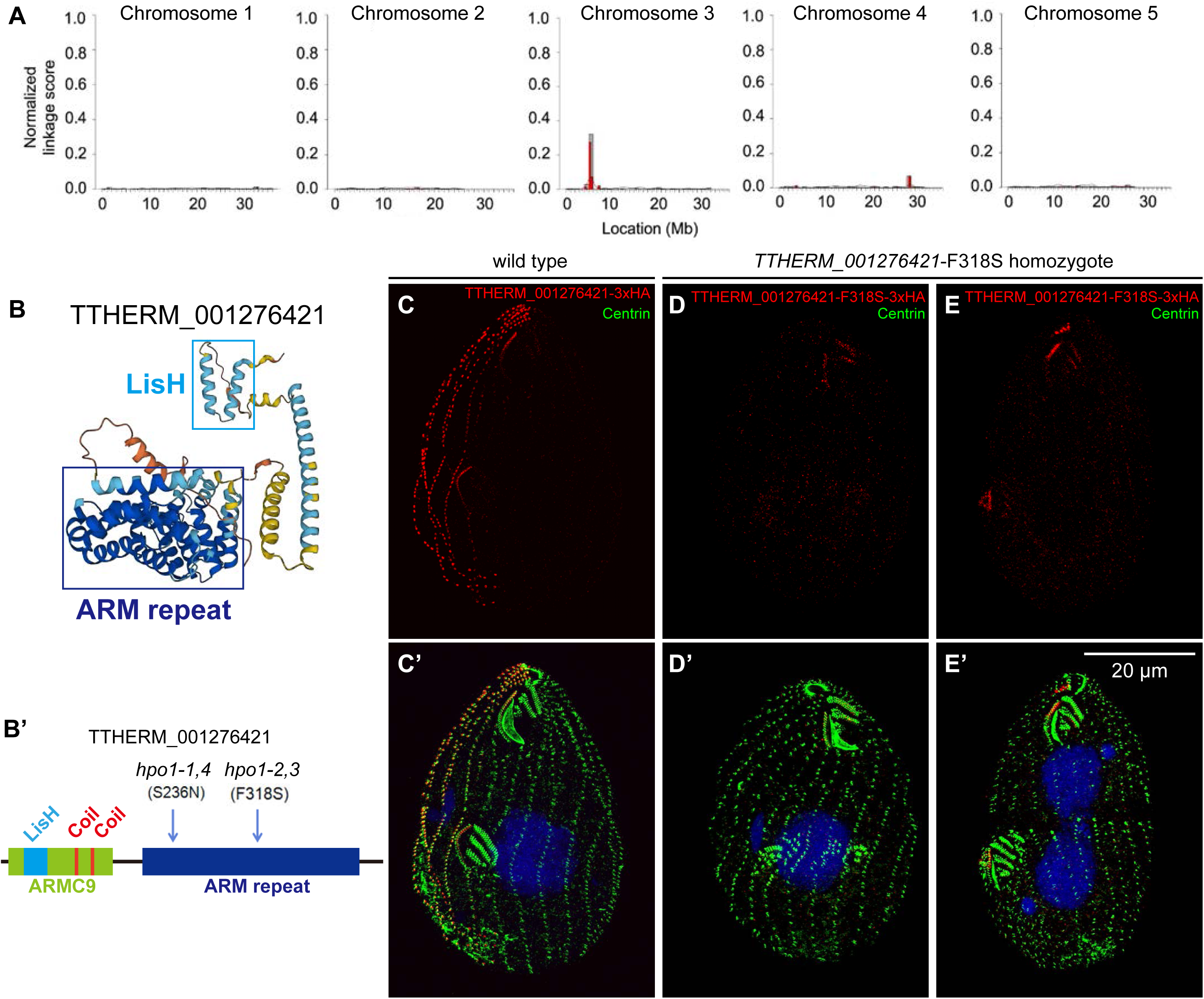
*hpo1* alleles map to *TTHERM_001276421* gene encoding an ARMC9-like protein. (A) Mapping of *hpo1-3* by the ACCA method. A linkage peak is present around 5 mB on the micronuclear chromosome 3. (B) The 3D protein structure of predicted TTHERM_001276421 protein generated by AlphaFold2. (B’) The domain organization of TTHERM_001276421 based on InterPro. The positions of two substitutions found in strains homozygous for the four *hpo1* alleles are marked. (C-E’) SR-SIM images show the cortical localization of either TTHERM_001276421-3xHA, a C-terminally tagged wild type protein (C,C’) or TTHERM_001276421-3xHA with the *hpo1-3*-linked substitution F318S (D-E’). The edited strain used was a homozygote expressing only the variant TTHERM_001276421-F318S protein. Note that expression of the F318S variant of TTHERM_001276421 phenocopies the cortical organization (multiple OPs) of *hpo1-3*. After growth at 38°C for 3 hours, the cells were labeled with the anti-HA antibody (red), 20H5 anti-centrin antibody (green), and DAPI (blue).

To test whether *TTHERM_001276421* is the locus of *hpo1* alleles, we used homologous DNA recombination to edit *TTHERM_001276421* to encode the *hpo1-2(3)* linked F318S variant with a C-terminal 3xHA tag. Homozygotes expressing TTHERM_001276421-F318S-3xHA had the phenotype similar to the one observed in the original *hpo1-3* mutants: multiple early-stage OPs forming near adjacent ventral rows (Fig. 3D,D’ compare to Fig. 3C,C’) or a single compound primordium (resulting from fusion of adjacent primordia) at a later cell division stage (Fig. 3E,E’). While the wild-type TTHERM_001276421-3xHA protein was enriched near anterior BBs along a subset of ciliary rows on the cell’s right side (Fig. 3C,C’) (see below for a detailed analysis of the localization pattern), the F318S variant was either not-detectable above the background (Fig. 3D-D’) or greatly diminished but still biased to the cell’s right side (Fig. 3E-E’ compare to 3C,C’). Thus, F318S may reduce the targeting of TTHERM_001276421 protein to the cell cortex or decrease its stability. Next, we created a germ-line based strain with a deletion of *TTHERM_001276421*. The *TTHERM_001276421-*KO homozygotes showed a phenotype similar to the original *hpo1* mutants with a subtle difference: the *knockout* cells had a higher number of OPs per cell as compared to the *hpo1-3* homozygotes (Fig. 2C,F,G). Furthermore, while in the *hpo1-3* homozygotes the OPs were predominantly shifted to the cell’s right side, in the *TTHERM_001276421*-KO cells the OPs were more frequently shifted to the cell’s left side at both 30 and 38°C (Fig. 2C,F compare to 2B,E, Fig. 2H). A leftward shift was reported to occur in the *hpo1-2* homozygotes grown at 39°C for 24 hr ^16^. Thus, the direction of the shift in the stomatogenic row position appears to depend on the degree of loss of function of Hpo1 and the original *hpo1* alleles are likely hypomorphs. Overall the phenotypes conferred by the engineered alleles are remarkably similar to the phenotypes observed in the original *hpo1* mutants ^16^ and therefore we concluded that *TTHERM_001276421* is the sought *HPO1* gene.

### Hpo1 forms a circumferential cortical gradient with a high point on the cell’s right side

During interphase, Hpo1-3xHA was enriched near the most anterior BBs of ∼6 somatic ciliary rows on the cell’s right side (Fig. 4A-A’’’). Along these rows, the levels of Hpo1-3xHA were high within the first most anterior 8-10 BBs and decreased along row length (Fig. 4A-A’’’). In addition, Hpo1 marked the BBs of the UM row in the OA (um in Fig. 4A’). On the ventral side, there was an abrupt drop in the Hpo1 level between row +1 and row 0 (Fig. 4A-A’). This seems important because the OP starts to form in the immediate proximity of BBs of row 0 (likely by nucleation from somatic BBs of row 0 serving as templates ^20^). In the early divider with a young OP, the Hpo1-3xHA pattern was unchanged, with a drop-off between rows +1 and 0 and the presence in UM (Fig. 4B-B’’’). In dividers with a prominent fission line, the pattern of Hpo1-3xHA was duplicated along the AP axis (Fig. 4C-C’’’). Namely, Hpo1-3xHA appeared along the anterior BBs of a subset of posterior ciliary half-rows on the right side of the cell, mirroring the positions of the pre-existing Hpo1-3xHA in the anterior hemi-cell (Fig. 4C-C’’’). Thus, the C pattern of Hpo1 is faithfully propagated during the cell cycle. Observations of live cells expressing GFP-Hpo1 revealed a pattern of Hpo1 distribution consistent with the data obtained in fixed cells (Fig. S2B,C). In addition to the localization at BBs, Hpo1 weakly localized at positions consistent with the microtubule bundles associated with the BBs (transverse and longitudinal microtubule bundles, Figure S2A,B,C). We did not detect displacements of GFP-Hpo1 foci.

**Figure 4.**
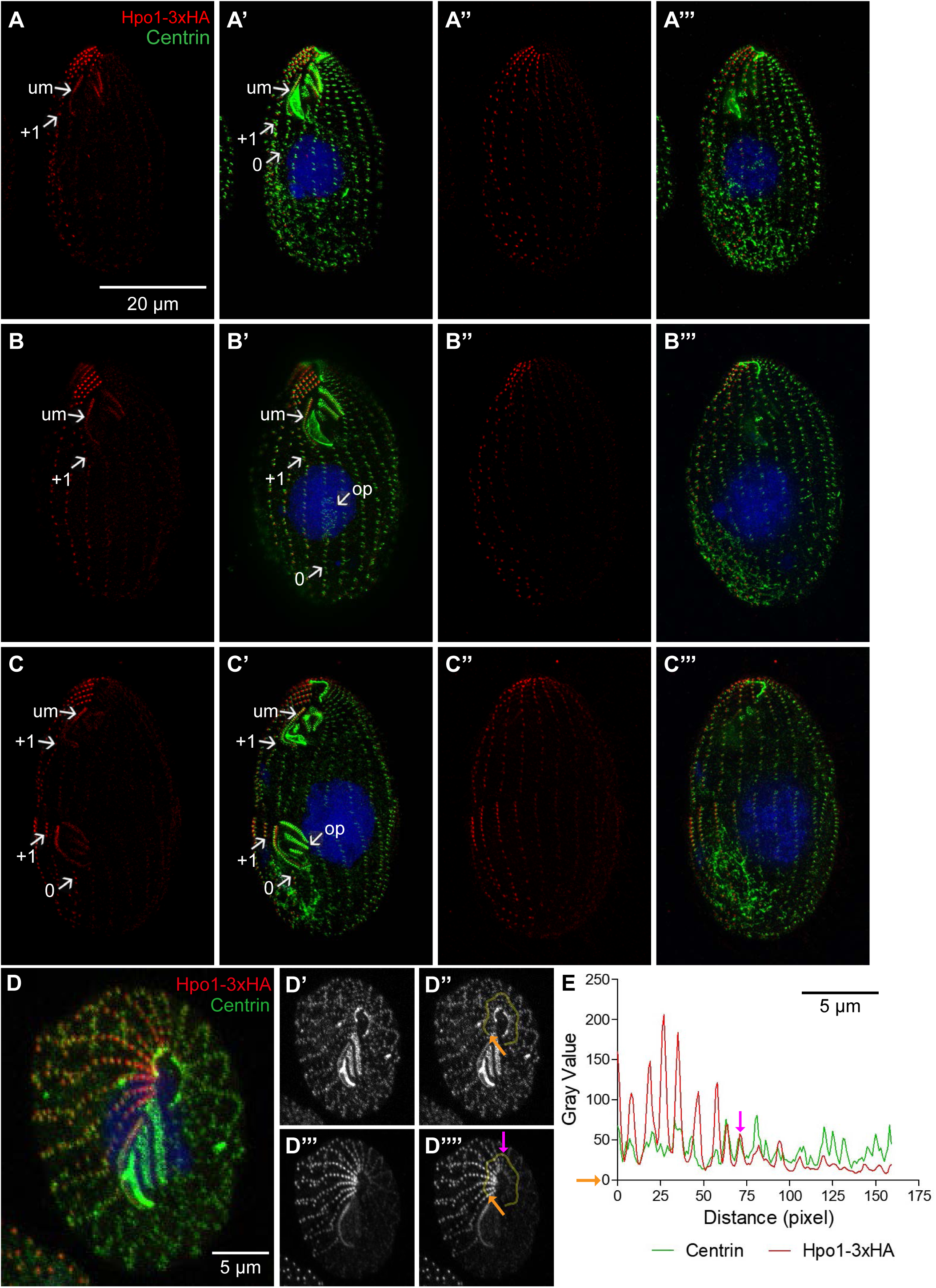
Hpo1 forms a bidirectional gradient with a high point on the cell’s right lateral side. (A-C’’’) SR-SIM images showing two sides of cells expressing Hpo1-3xHA during the cell cycle. Cells were labeled with the anti-HA (red) and 20H5 anti-centrin (green) antibodies, and DAPI (blue). (D-D’’’’) Confocal images of an apical fragment of a cell expressing Hpo1-3xHA. The cell was labeled with anti-HA (red) and 20H5 anti-centrin (green) antibodies, and DAPI (blue). Single channel gray scale images for centrin (D’,D’’) and Hpo1-3xHA (D’’’,D’’’’) were generated using the confocal image shown in D. (E) The intensity plots document the gray values measured across the region marked by the yellow lines shown in panels D’’ and D’’’’. The orange arrows indicate where the measurements were started, and the purple arrows indicate the position of the dorsal discontinuity. Abbreviations: um, undulating membrane row within the OA; op, oral primordium. The numbers mark ventral row positions.

A close inspection of the levels of Hpo1-3xHA around the cell’s circumference revealed a second drop-off on the cell’s dorsal side (Fig. 4A”-A”’, B”-B”’, C”-C”’). In the cells stained for fenestrin, a marker of CVPs ^21,22,^ the drop-off was apparent between the second and third row past the CVP row counting clockwise (Fig. 5C’). To map the position of the dorsal Hpo1 drop-off more accurately, we imaged the cell’s apical region in cell fragments that offered a “polar” view of the circumference (Fig. 4D-E). The centrin signal was nearly uniform around the cell circumference (Fig. 4D-E). The Hpo1-3xHA signal was highest at row +4 and decreased to almost the background level in two steps: between row +5/+6 and +7/+8. (Fig. 4D,D’’’,D’’’’,E).

**Figure 5.**
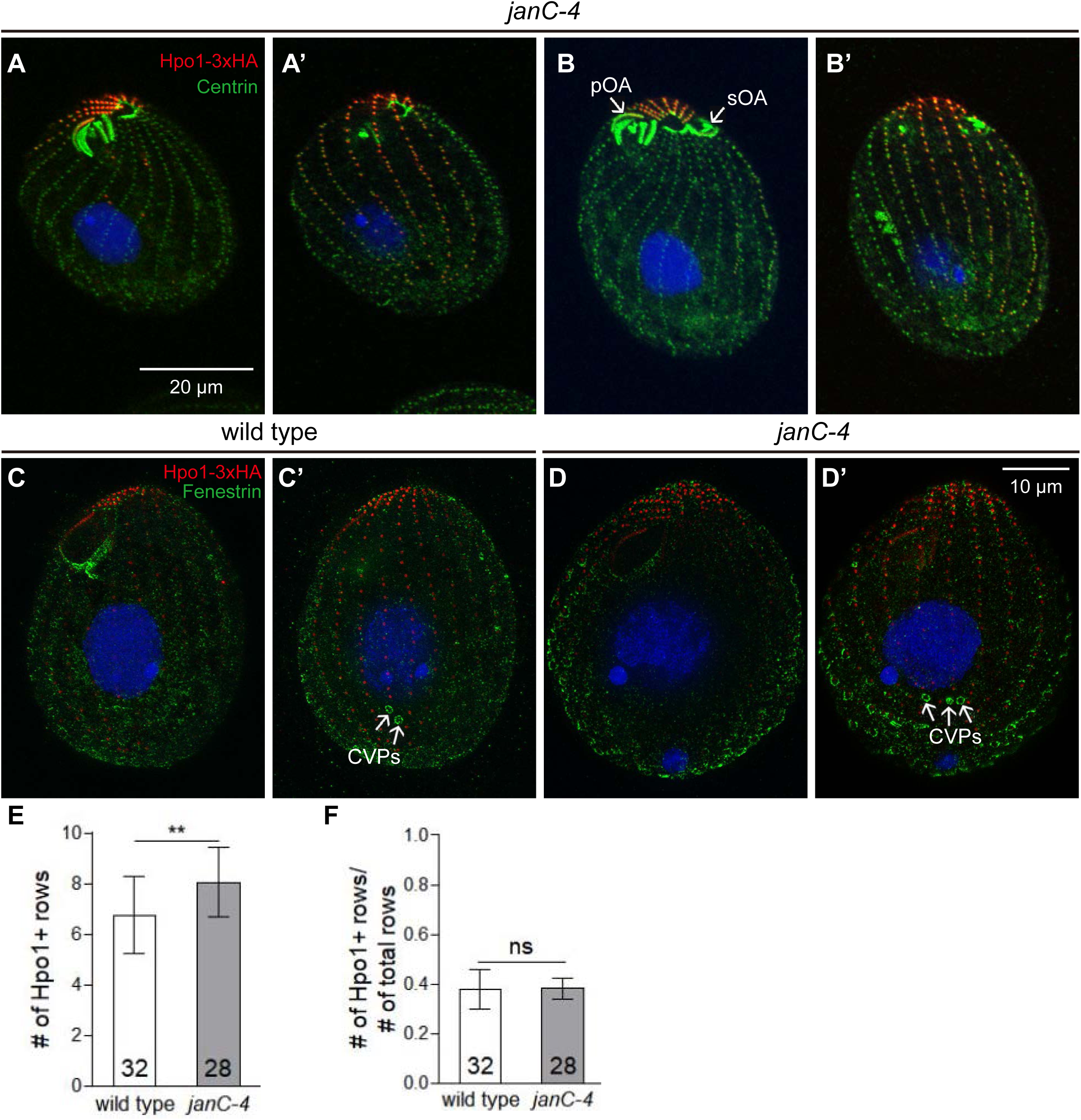
The dorsal discontinuity in the Hpo1 gradient marks a cryptic position for oral development expressed in the *Janus* mutant. (A-B’) Pairs of confocal images showing two sides of *janC-4 homozygote* cells that have either a single OA (A’A’) or two OAs (B-B’). Cells were labeled with the anti-HA (red) and 20H5 anti-centrin (green) antibodies, and DAPI (blue). (C-D’) SR-SIM image pairs of cells that are either wild-type (C,C’) or *janC-4* homozygotes (D,D’) and express Hpo1-3xHA. The cells were decorated with the anti-HA (red), anti-fenestrin (green) antibodies, and DAPI (blue). (E,F) Graphs reveal an increase in the number of rows with enriched Hpo1-3xHA in the *janC-4* background as compared to the wild type (E) but the fraction of the circumference occupied by high Hpo1-3xHA remains unchanged when the increase in the total number of rows is taken into account (F). ns: not significant, Stars indicate statistically significant (*: P < 0.05, **: P < 0.01, and ***: P < 0.001). Abbreviations: pOA, primary oral apparatus; sOA, secondary (dorsal and usually defective) oral apparatus; CVP, contractile vacuole pore.

Overexpression of Hpo1-3xHA using the cadmium-dependent promoter *MTT1*^23^ resulted in strong accumulation in the cell body. However, the right-side- and anterior-biased localization of Hpo1 was still visible and cortical positioning was not disturbed (Fig. S3). Likely, the biased localization of Hpo1 requires its binding to another spatially-biased cortical factor, and Hpo1, while required, is not rate-limiting for precise positioning of organelle development.

### The dorsal Hpo1 drop-off marks the position where an extra oral apparatus assembles in the *janus* mutant

In the *janus* mutants (*janA*, *janB* and *janC* allelic groups), ventral structures are abnormally duplicated on the dorsal cell surface ^24-27^. The dorsal (secondary) OA is underdeveloped and often inverted in its left-right internal polarity (sOA in Fig. 5B; Fig. S4A). The *janus* phenotype was interpreted as a global mirror-image pattern duplication ^24,25,27^. In the *janC-4* homozygotes the distribution of Hpo1-3xHA was similar to that in the wild type, including an enrichment of Hpo1 on the cell’s right side (Fig. 5A-B’ compare to Fig. 4). The average number of rows with high Hpo1-3xHA (rows between the two drop-off positions) was elevated from 7 in the wild type to 8 in the *janC-4* homozygotes (Fig. 5E), but *janC-4* cells had more total rows per cell and therefore the ratio of Hpo1-enriched rows to the total cell circumference was unchanged (Fig. 5F). The expression of the sOA in *janus* mutants is partially penetrant ^25-27^. Both *janC-4* mutant cells with and without a sOA had a similar number of Hpo1-enriched rows (Fig. 5B,B’ compare to 5A,A’). Strikingly, in cells with a fully expressed *Janus* phenotype, the sOA was located at the position of the dorsal discontinuity of Hpo1 (next to row +7 in Fig. 5B,B’; Fig. S4). Thus, in the *janus* background, Hpo1 is a bilateral marker for oral development in the *janus* background.

The *janus* mutants also have an increased number of CVPs that often are arranged as two sets separated by 1-2 rows lacking these organelles ^24,25^. Thus, there is a correlation between the widening of the Hpo1-enriched domain and the increased number of CVPs in the *janC-4* homozygotes. The mid-point of the CVP domain was located along or close to the row with the peak level of Hpo1 in both the wild type (Fig. 5C’) and *janC-4* cells (Fig. 5D’).

To summarize, OA development occurs at positions where Hpo1 levels suddenly drop off while CVPs form at the peak level of the Hpo1 circumferential gradient.

### Hpo1 acts bidirectionally to restrict circumferential positions competent for oral development

Based on its circumferential distribution pattern in the *janC-1* homozygotes, Hpo1 may act bidirectionally to exclude OPs from the region of its enrichment on the cell’s right lateral side. To test this idea, we analyzed double mutant homozygotes, *janC-1;hpo1-3.* The double mutants presented a highly penetrant phenotype, two mature OAs were located side by side near the anterior cell end, sometimes separated by a gap of one or more ciliary rows (Fig. 6D compare to 6A-C). While such cells were rare in the single *janC-1* or *hpo1-3* mutants, the frequency reached 70% in the double mutants (Fig. 6I,J). When such twin OAs were physically adjacent to one another with no intervening gaps, they often appeared to integrate into a single, compound organelle (Fig. 6E). It appears that under Hpo1 deficiency, the two OAs exhibit “cortical slippage” (caused by ongoing shifts in the positions of both OPs) that eventually result in a collision of the formerly separate OPs within the *janC* cortex (a situation predicted by ^14^) (Fig. 6K). Curiously, the direction of cortical slippage appears reversed for the two oral domains: OP slippage appears to occur to the cell right of the primary oral meridian, and to the cell left of the secondary meridian (Fig. 6K). Given its high frequency, the “twin OA” configuration appears relatively stable and is likely propagated during division. Occasionally we observed dividing cells seemingly in the course of “cortical slippage” where the distance between the OAs decreased within one generation; the dividing cells in Fig. 6F-H have two old OAs still separated by a gap, and multiple immediately adjacent OPs. Overall, these data suggest that Hpo1 acts to separate the regions competent for oral development, one of which (dorsal) is typically repressed in the wild type through action of the *Janus* gene products. These data correlate the bilateral gradient pattern of Hpo1 (described above) with its bilateral (OP excluding) activity.

**Figure 6.**
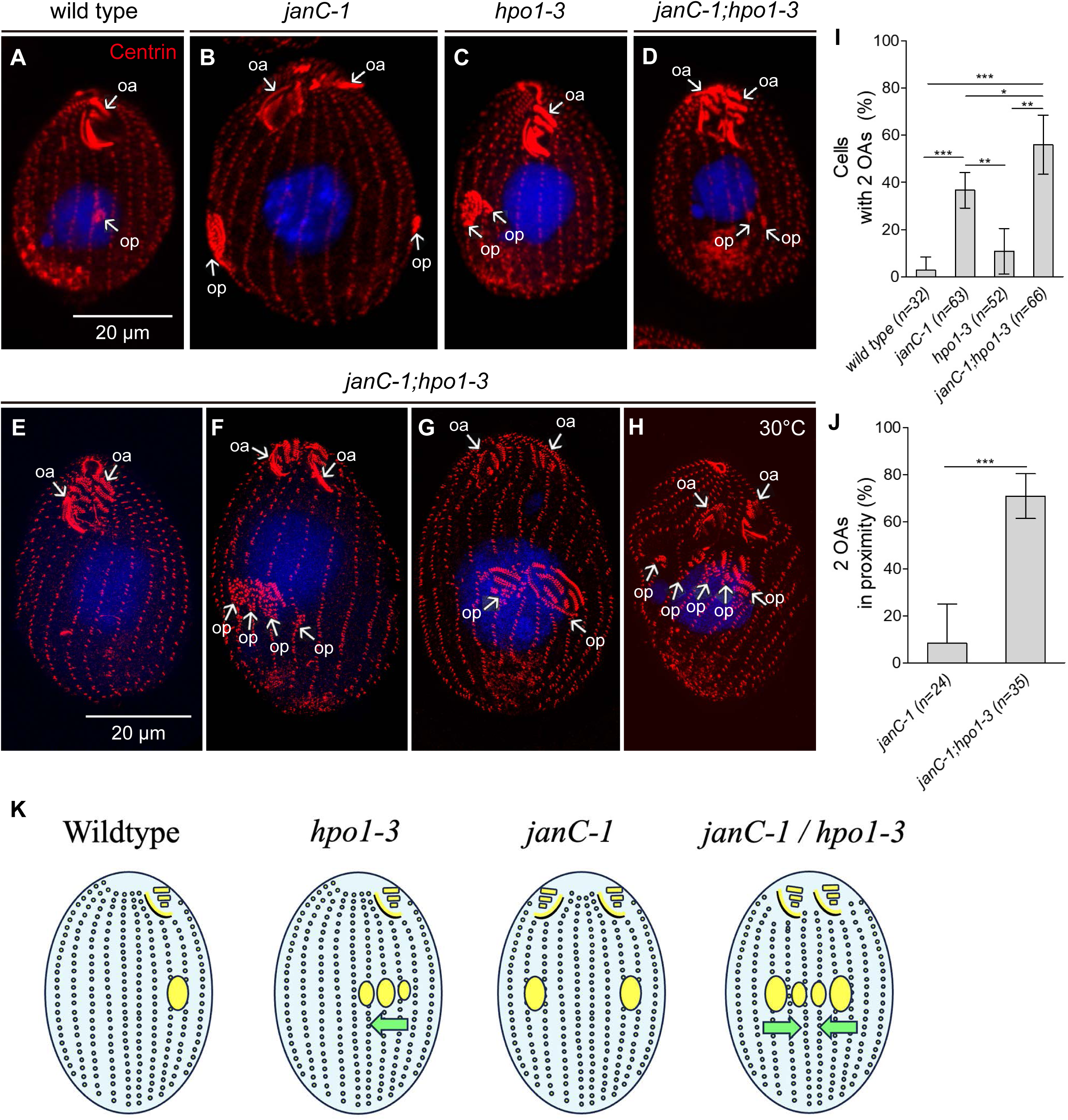
Hpo1 acts bidirectionally to separate the two oral apparatuses in the *janus* mutant. Confocal (A-D) and SR-SIM (E-H) images cells with indicated genotypes labeled with the 20H5 anti-centrin antibody (red), and DAPI (blue). Cell shown in all panels except H were incubated for 4 hours at 38°C to enhance the *hpo1-3* phenotype. (I,J) The graphs quantify the frequencies of cells two mature OAs (I), and two OAs in close proximity (J). The bars represent the means ± SD. Two-tailed unpaired t-test was executed for statistical analysis. ns: not significant. Stars indicate statistically significant (*: P < 0.05, **: P < 0.01, and ***: P < 0.001). (K) The drawing summarizes the outcome of the genetic interaction between *janC-1* and *hpo1-3*. The yellow ovals are OPs. The green arrows shows the proposed predominant direction of shifts in the OP positions. Abbreviations: pOA primary OA; sOA secondary OA; op, oral primordium.

### Hpo1 interacts with Bcd1, a left side-enriched OP positioning factor

Bcd1 is a conserved Beige-BEACH domain protein ^7^ whose loss of function in *Tetrahymena* confers a phenotype superficially similar to that of *hpo1*: multiple adjacent oral primordia ^15^. However, while the *hpo1* alleles reduce the number of CVPs (Fig. S1B,C,H), *bcd1* alleles increase the number of CVPs per cell ^7,15^ and (Fig. S1F,H). Also, while Hpo1 is enriched on the cells’ right side, Bcd1 is enriched on the cell’s left side where it appears also to form a bidirectional circumferential gradient when viewed toward the apical cell end (See Fig. 5H in ^7^, and Fig. S5A-B”’). The AP distribution of Bcd1 protein resembles that of Hpo1, high near the most anterior row segments and fading away toward the posterior row ends (^7^ and Fig. S5A-B”’).

The similarity of *hpo1* and *bcd1* OP phenotypes and the enrichment of Hpo1 and Bcd1 proteins on opposite sides of the stomatogenic ciliary row suggest that the two proteins act collectively to restrict the OP formation from spreading to either the cell’s right (Hpo1) or cell’s left (Bcd1) of this prominent cortical landmark. To look for potential interactions between Hpo1 and Bcd1, we first examined whether a deficiency of Bcd1 (*bcd1-2*) affects the pattern of Hpo1-3xHA. In the *bcd1-2* homozygotes, the average number of Hpo1-enriched rows (rows between the discontinuities) was increased to 9 (Fig. S6F) but the total number of rows per cell also increased and therefore the Hpo1-enriched domain was proportionally unchanged (Fig. S6G). As in the wildtype, Hpo1 signal showed a dramatic drop-off in concentration both ventrally (near the primary oral meridian) and dorsally (Fig. 7B-B” compare to 7A-A”; Fig. 7C,E). However, the *bcd1-2* background consistently altered the AP distribution of Hpo1-3xHA. In wild type cells, the anterior-posterior Hpo1-3xHA gradient is steep, dropping to half signal intensity within ∼10 BBs of the anterior cell end. In *bcd1-2* homozygotes, the AP gradient of Hpo1-3xHA remained high, only dropping to 75% signal intensity (Fig. 7B-B” compare to 7A-A”; Fig. 7D). This gradient-altering effect was observed both on the cell’s ventral and dorsal side (Fig. S6).

**Figure 7.**
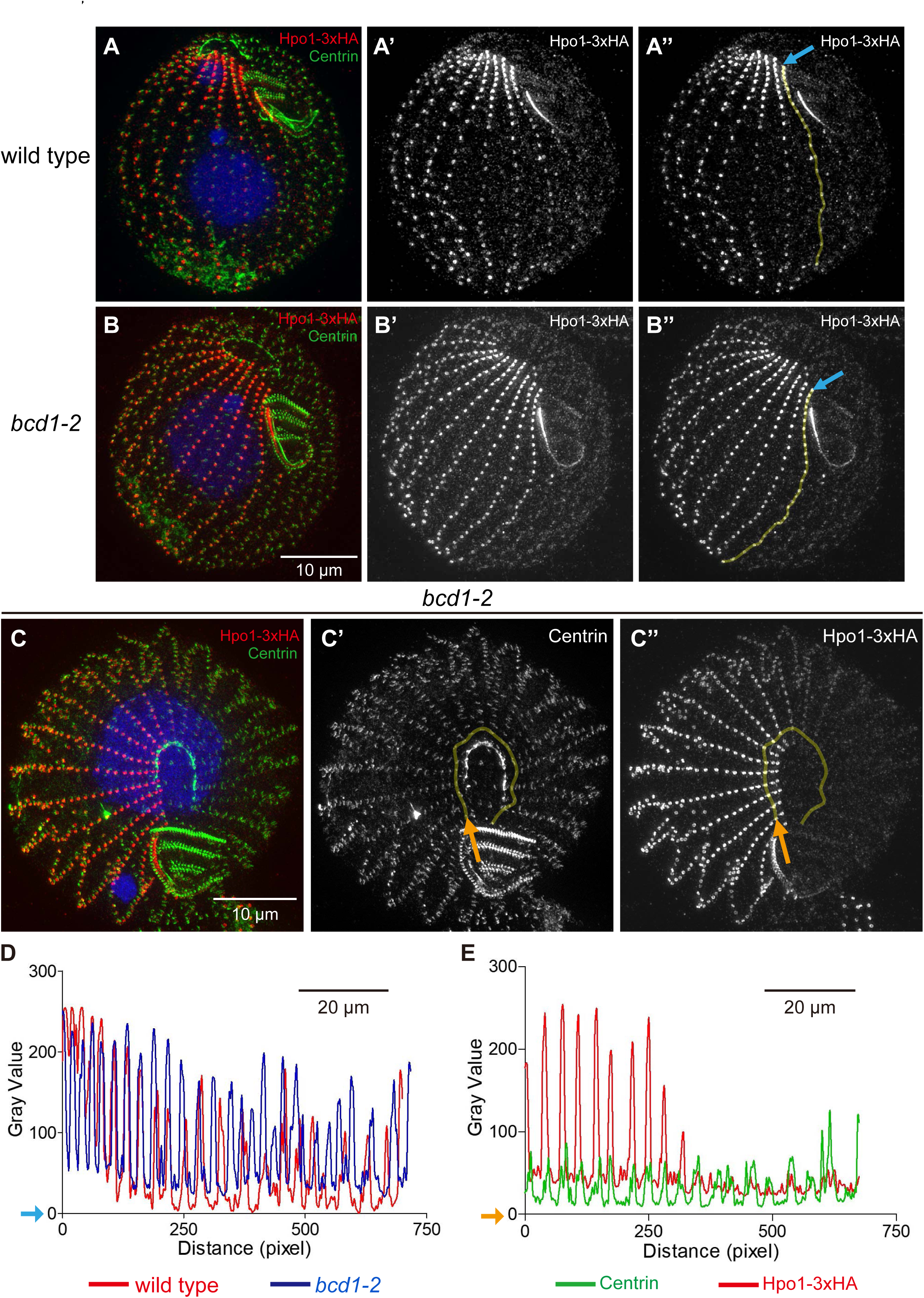
A Bcd1 deficiency changes the A/P distribution of Hpo1. (A-B’’) SR-SIM images of Hpo1-3xHA-expressing cells that are either otherwise wild-type (A-A’’) or *bcd1-2* homozygotes (B-B’’). The cells were labeled with the anti-HA (red), 20H5 anti-centrin (green) antibodies, and DAPI (blue). The grey scale images are the single channel signals of Hpo1-3xHA. The yellow lines (A’’ and B’’) cover the region whose gray values were measured along the A/P axis, and the light blue arrow indicates the initial point of measurement. The corresponding A/P grey value intensity plots for Hpo1-3xHA in the two genetic backgrounds are shown in the graph in panel D. (C-C’’) An SR-SIM image of the cell’s apical region from a cell expressing Hpo1-3xHA (red) in the *bcd1-2* background, also stained with anti-centrin (green) antibody. The single channel grey scale images for centrin (C’) and Hpo1-3xHA (C’’) were used for measurements of the circumferential signal intensities. The measured regions are marked by the yellow lines and the orange arrows orient the measurements. The resulting signal intensity plots are shown in the graph in panel E.

Next, we focused on whether the C distribution of Hpo1 is affected by *bcd1-2*. In wild-type cells, the ventral Hpo1 drop-off occurs between row +1 (contrast row) and row 0. In the dividing *bcd1-2* cells in which the OP fields were shifted laterally, the contrast row position was consistently shifted to the row on the right side of the right-most OP. For example, in the dividing cell shown in Fig. 8B-B’, there are two early OPs at positions 0 and +1 and the contrast row is located to the right of the OP pair at row +2. While we have not observed clear cases of a shifted contrast row in interphase cells, there were instances of apparent ambiguity in position of the contrast row. In the cell shown in Fig. 8A-A’, within the anterior 1/3 of the cell the contrast row is +1. However, in the posterior 2/3 of the cell the contrast row is +2. This cell has 3 postoral rows, a feature common to both *bcd1* and *hpo1* mutants that correlates with the increased OA width. However, another interphase *bcd1-2* cell shows the same “ambiguous contrast row” phenotype in the presence of two postoral rows (Fig. 8D-D’). Taken together, these data reveal that the Bcd1 deficiency modifies both the AP and C pattern of Hpo1. These observations suggest that there is a cross-talk between Hpo1 and Bcd1, possibly as a boundary interaction between the right and left ventral cortical region.

**Figure 8.**
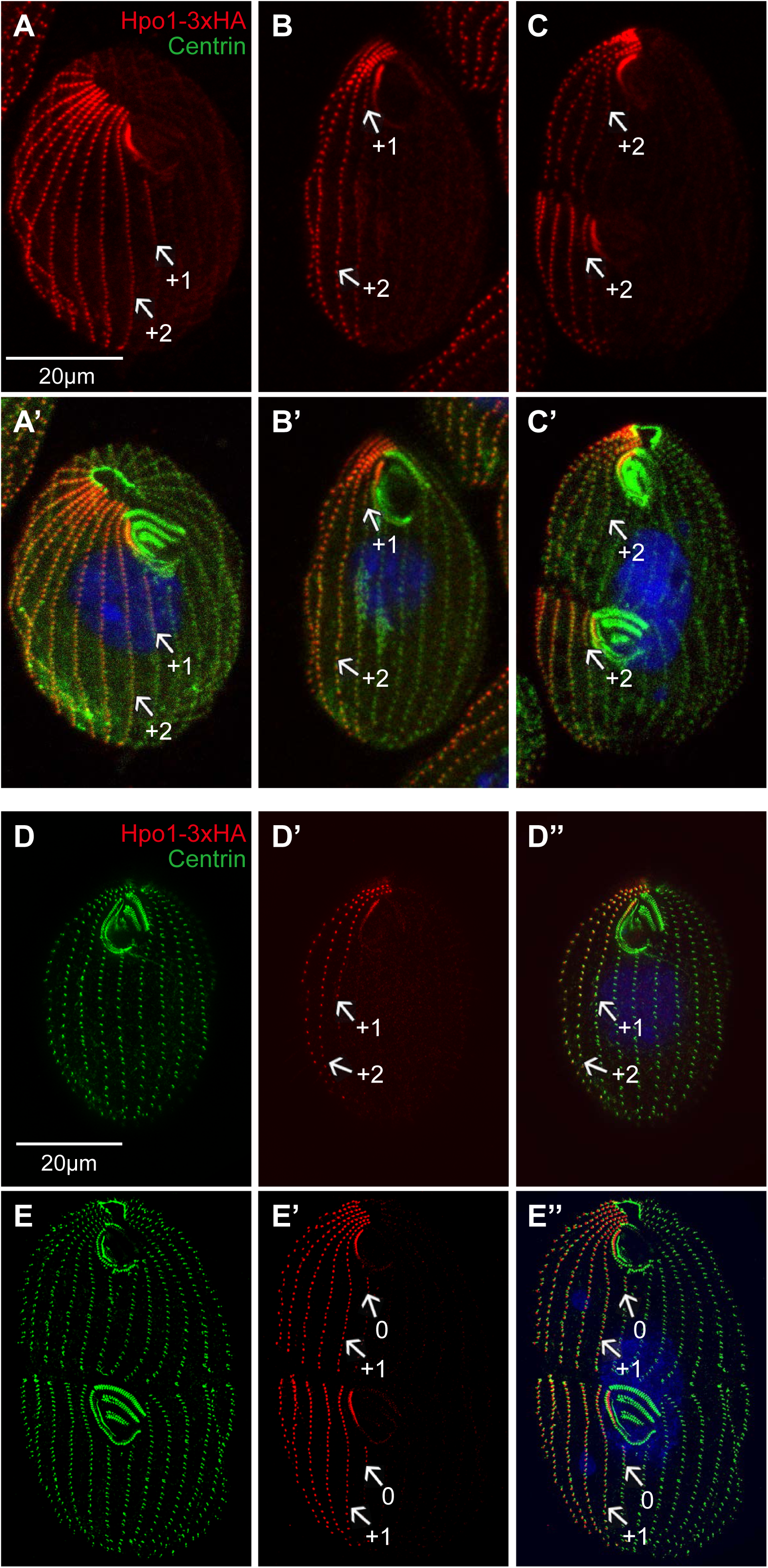
Bcd1 influences the circumferential pattern of Hpo1-3xHA. Confocal (A-C’) and SR-SIM (D-E’’) images of cells expressing Hpo1-3xHA that are homozygous for *bcd1-2*. After overnight incubation at 30°C, the cells were labeled with the anti-HA (red), 20H5 anti-centrin (green) antibodies, and DAPI (blue). The numbers mark the ventral row positions.

No obvious effect of expression of *hpo1-3* on the distribution of Bcd1-GFP was observed but the signal of Bcd1-GFP was weak even in the wild-type background making the evaluation difficult (Fig. S5C-D”’ compare to Fig. S5A-B”’).

We next asked whether there is a functional interaction between Hpo1 and Bcd1 by analyzing the phenotypes of double mutants homozygous for *bcd1-2* and *hpo1-KO*. Surprisingly, the *bcd1-2;hpo1-KO* homozygotes were not viable. To test further for synthetic lethality, we attempted to rescue the progeny of mating (double mutant) heterokaryons by biolistic introduction of a transgene mediating expression of Hpo1-3xHA in an unrelated locus. A total of about 5 x 10^6^ mating heterokaryon cells were either subjected to biolistic bombardment with a Hpo1-3xHA transgene or mock-transformed without plasmid DNA and selected with paromomycin (conferred by the *neo* gene embedded in the disrupted *HPO1* gene). Eighteen drug-resistant clones were isolated from the population bombarded with the Hpo1-3xHA plasmid and one clone was isolated from the mock-transformed population. Six clones from the plasmid bombarded cells were analyzed by immunofluorescence and all were positive for Hpo1-HA (Fig. S7B-C’). The single clone selected in the mock-transformed population was negative for Hpo1-HA as expected (Fig. S7A,A’). A PCR amplification detected a wild-type Hpo1 sequence corresponding to the portion that was deleted in the *hpo1*-KO allele in the “escapee” clone (Fig. S7D,D’). Likely, in this single clone there was a transfer of the wild-type *HPO1* gene from the parental macronucleus to the newly developing macronucleus that rescued the lethality. These observations confirm that the loss of function of Bcd1 and Hpo1 is lethal and 100% penetrant (n=5 x 10^6^). Next, we evaluated how the cortical pattern changes during the transition from the wild-type to double mutant phenotype in the mass-selected progeny of mating heterokaryons. To this end, heterokaryons expressing complementary mating types were allowed to mate, refed and selected with paromomycin to kill the non-mating (phenotypically wild-type) parental cells. After 24 hr of drug selection, the double mutant progeny cells looked nearly normal except for the frequent OAs with abnormally wide oral M rows (white arrows in Fig. 9A). After 30-46 hr, dividing cells frequently had severely posteriorly displaced primordia (yellow arrows in Fig. 9B,E,F,G). In the cells with a posteriorly shifted OP, the division boundary position was variable, either also shifted to the posterior (Fig. 9F) or equatorial (Fig. 9E) or shifted anteriorly (Fig. 9G). Cells with a posteriorly shifted OP and an anteriorly shifted division plane may produce a posterior daughter cell with mature OA near the posterior cell end (Fig. 9H). At ∼46 hr the cells were uniformly arrested in cell proliferation and had an abnormally elongated and curved morphology with OAs that had curved (pink arrows in Fig. 9C,I) or fragmented M rows (green arrows Fig. 9C). The somatic ciliary rows appeared excessively long, and sometimes twisted (Fig. 9D,H,I). SR-SIM imaging revealed additional patterning defects including OAs that lacked the UM (Fig. 9G,I) and OAs with M rows arranged with orthogonal orientations (small arrows in Fig. 9J,K). In addition, there were defects in the organization of the buccal cavity including a missing or fragmented ribbed wall of microtubules (Fig. 9G,I,J,K). The *bcd1-2*;*hpo1*-KO heterokaryon progeny did not grow on the specialized culture medium (MEPP) that supports proliferation of mutants lacking a functional oral apparatus ^28^. Thus, the lethality is not solely caused by a lack of a functional oral apparatus. We conclude either Hpo1 or Bcd1 are required for survival and both contribute to patterning functions that extend beyond positioning of the OP, including shaping the internal organization of the OA and controlling the longitudinal expansion of the somatic ciliary rows.

**Figure 9.**
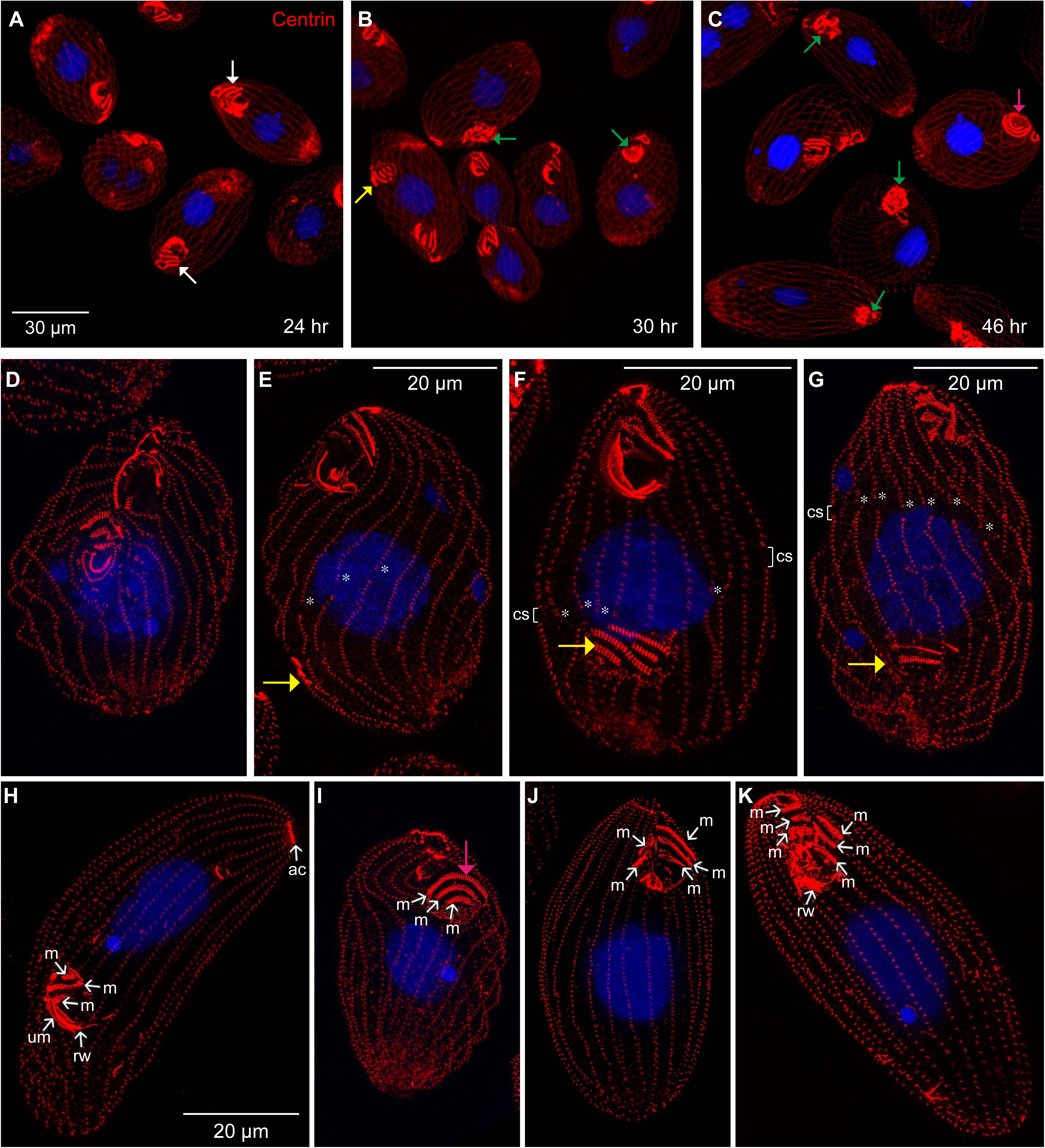
A loss of both the left-side enriched Bcd1 and the right-side enriched Hpo1 severely disturbs cortical patterning. Confocal (A-C) and SR-SIM (D-K) images of *bcd1-2*;*hpo1*-KO double homozygotes obtained as progeny of mating heterokaryons. The times shown in panels A-C refer to the period after refeeding of the mating heterokaryons (after 18 hr of mating at 30°C). Cells were labeled with the 20H5 anti-centrin antibody (red), and DAPI (blue). Abbreviations: m, M oral row; um, UM oral row; rw, ribbed wall of microtubules; ac, apical crown; cs, cortical subdivision. The stars show the gaps in the longitudinal rows that indicate the position of the “cortical subdivision” (a set of gaps in rows at the division plane). Arrows of multiple colors indicate types of cortical defects: white, OAs with long oral rows; yellow, OP displaced posteriorly; green, OA with disorganized rows; pink, OA with circular rows.

## Discussion

Ciliates are well suited for studies on intracellular pattern formation but remain relatively unexplored. In *Tetrahymena*, mutations in several loci selectively affect organelle positioning along either the AP or C axes (reviewed in ^1,14^), suggesting that pathways operating on each of the two orthogonal polarity axes have a degree of independence. Several highly conserved kinases and kinase-binding proteins, including components of Hippo signaling, mediate organelle positioning along the AP axis ^29-34^. The emerging view is that on the AP axis, organelle locations are determined by proteins that mark cortical domains within which new structures are not permitted to form (reviewed in ^1^). These inhibitory cortical factors are either permanent (Elo1/Lats ^32^) or appear shortly before the structures form (CdaI/Mst ^31^, CdaA/cyclin E ^29^). Here we made an advance toward understanding the far less explored mechanism of patterning around the cell circumference in *Tetrahymena*, by identifying the *HPO1* gene. The *hpo1* alleles disturb circumferential patterning by permitting oral development outside of the standard row 0. Over generations, the ongoing shifts in the positions of oral meridians (cortical slippage) may even produce mutant cells that have CVPs on the left cell’s side, and therefore have an inverted overall “handedness” ^16^.

Hpo1 is a ciliate phylum-specific BB-associated ARMC9-like protein. In *Tetrahymena* and mammalian cells ARMC9 localizes to BBs and the tips of cilia ^17,18,35^. Mutations in ARMC9 cause Joubert syndrome, a neuro-developmental ciliopathy ^18,19^. Likely, Hpo1 evolved by neofunctionalization of the ancestral ARMC9, following a gene duplication that occurred before the emergence of diverse ciliate subclasses. We recently identified another conserved ciliary protein, a Fused/Stk36 kinase CdaH, as a key regulator of positioning on the AP axis ^34^. Ciliates belong to the protist clade of Alveolata that also includes dinoflagellates and apicomplexans. Among the alveolate protists, only ciliates assemble multi-ciliated arrays. Thus, in the course of emergence of ciliates from an ancestral biciliated alveolate ^36-38^, multiple ciliary proteins could have gained functions in the formation and patterning of ciliary rows.

Hpo1 forms concentration gradients along both cell polarity axes. Remarkably, on the circumferential axis, Hpo1 forms a bilateral concentration gradient with the high point on the cell’s lateral right side and sharp drop-offs along the longitudes permissible of oral development. The pattern of Hpo1 appears invariable during the cell cycle. Hpo1 is reminiscent of Elo1, a Lats kinase that marks the posterior cell region and acts to prevent the OP from forming too close to the posterior cell end ^32^. Thus, the positioning on both orthogonal polarity axes involves gradient-forming proteins that may function as pre-existing markers that define positions at which organelles assemble.

Important early insights into the mechanism of circumferential patterning were obtained using the giant ciliate Stentor coeruleus. *Stentor* species have formidable healing and regenerative capabilities. Fragments of *Stentor* lacking most of the cortex can regenerate a complete pattern and recover the ability to grow and divide, which argues that pre-existing cortical organelles are not an essential source of positional information for forming structures ^39-41^. Importantly, in *Stentor*, the circumferential surface differentiation is apparent along the entire cell length. Unlike in *Tetrahymena*, in *Stentor*, the width of the inter-row spaces (called stripes) gradually changes around the cell circumference. The narrowest stripes are located on the right side of the ventral surface and the stripe width gradually increases clockwise until the widest stripes meet the narrowest stripes within the ventral region named the “locus of stripe contrast” (LSC) ^42,43^. OP develops at the LSC, within the narrow stripe zone and close to the margin of the wide stripe zone ^43-45^. Remarkably, supernumerary OPs can be induced by cortical grafts that juxtapose a narrower stripe cortex with a wider stripe cortex, even if the boundaries of the graft would normally be far from the sites of cortical development ^46^. Thus, oral development occurs at locations that are characterized by a “structural contrast”. In *Tetrahymena*, Hpo1 forms a “molecular contrast” as the sharp level discontinuity between row +1 and the stomatogenic row 0. Topologically, the Hpo1-enriched region in *Tetrahymena* corresponds to the narrow stripe region in *Stentor*. Taken together, our observations and the grafting studies in *Stentor* suggest that the ventral “contrast region” contains both activating and inhibitory influences that control oral development. We speculate that the right lateral region, where Hpo1 is concentrated, is also a source of an unknown “oral activator” that is inhibited by Hpo1 (green gradient in Fig. 1C). Hpo1 and the oral activator gradients may overlap for most of the length except at the positions where the Hpo1 levels drop, thus creating conditions permissible for oral development (Fig. 1C). Consequently, (in cells having a normal Janus activity, see below) the OP can form at row 0.

By analogy to the emerging principle of positioning on the AP axis (where pattern regulators mark cortical domains where organelle assembly is inhibited), the primary activity of Hpo1 could be an exclusion of oral development from the cell’s right lateral side. Indeed in all original *hpo1* mutants (expressing alleles that are likely hypomorphs), the extra primordia form more frequently to the right of row 0 (^16^ and this study). Unexpectedly however, in cells with a knockout of *HPO1,* there was an increase in the frequency of extra primordia to the left of row 0. The *hpo1-2* mutants also occasionally form primordia on the left side and that the frequency of left-shifted OPs increases at 39°C (^16^ and this study). We speculate that in addition to the exclusionary activity to the right of row zero, Hpo1 stimulates the exclusionary activity to the left of row 0, which requires Bcd1. Both Hpo1 and Bcd1 mutants assemble extra oral primordia on either side of row 0 (^15,16^ and this study). It is tempting to speculate that Bcd1 and Hpo1 have OP excluding activity in the areas of their enrichment (Hpo1 on the right and Bc1 on the left) and enhance each other’s exclusionary activity as a boundary effect across row 0 (Fig. 1C).

We show that Hpo1 is organized as a bilateral gradient with drop-offs on the dorsal and ventral side. Both drop-off positions mark sites for oral development in the *janC* homozygotes. *janA* and *janB* alleles confer the same *janus* phenotype as the *janC* alleles used here ^47-49^. The JanA gene product was recently identified as a Polo kinase that localizes to the left-to-dorsal circumferential region, with a large overlap with Bcd1 and a partial overlap with Hpo1 on the dorsal side ^8^ (see Fig. 1C). Cole and colleagues propose that JanA represses the expression of ventral features on the dorsal side by interfering with a right-side enriched oral activator ^8^. We show here that in the double mutant expressing *janC-1*, a loss of Hpo1 reduces the distance between the primary and the secondary OAs. This observation suggests that Hpo1 has bilateral activity, excluding oral development from both dorsal and ventral margins of its gradient. Hpo1 exclusion of dorsal oral development is masked by the over-arching Janus suppression activity. The Janus gene products may either act to modify the parameters of the Hpo1 gradient (e.g. by reducing its steepness at the dorsal drop-off position) or suppress downstream components required for expression of oral structures. While in cells having wild-type Janus gene products, the right-sided influence of Hpo1 on the OP position is not relevant, on the dorsal side Hpo1 may engage in other positional activities, including its potential dorsal interaction with Bcd1 that may control the number of CVPs (see below).

Intriguingly, all gene products studied here in addition to their effects on oral development, also affect the number of CVPs. While all alleles used here either null or hypomorphs, *hpo1* alleles decrease while the *jan* and *bcd1* alleles increase the number of CVPs, respectively. While during cell division, the OP forms before the CVPs, it is unlikely that either the number or positions of OPs affect the number of CVPs based on the phenotypes of *hpo1* and *bcd1* mutants that affect oral development in a similar way but have opposite effects on the number of CVPs. We observed that CVPs are located at the posterior ends of rows whose anterior ends are at or near the high point of the Hpo1 circumferential gradient. Furthermore, we show that a loss of either JanC or Bcd1 expands the Hpo1-enriched domain. Both Bcd1 ^7^ and JanA are located in the left dorsal region ^8^ and partly overlap with Hpo1 and could act on the CVP rows indirectly by regulating the parameters of the Hpo1 gradient.

Unexpectedly, by combining null alleles *hpo1*-KO and *bcd1-2,* we uncovered that either the right-side-enriched Hpo1 or left-side-enriched Bcd1 are required for *Tetrahymena* survival. The double mutants lacking both Hpo1 and Bcd1 arrest in the cell cycle with a grossly misshaped cortical pattern. One of the prevailing phenotypes in the double mutants are frequent posterior shifts in the OP position. This phenotype was earlier seen in the *hpo1-2* mutants exposed to a higher temperature for a prolonged period ^16^ and occasionally in the *bcd1-2* mutants (see Fig. 4A in ^15^). Frankel and colleagues suggested that Hpo1 may be a shared component of pathways that operate on the C and AP axes ^16^. We note however that all C patterning factors studied to date (Bcd1, JanA and Hpo1) in addition to the circumferential distribution bias, also show a graded distribution along the AP axis (^7,8,^ and this paper). Thus, there could be a deeper relationship between the C and AP positioning pathways.

To our knowledge, we are first to document essential interactions between the right and left side-enriched pattern regulators. These observations bring to mind some fascinating cortical graft experiments performed on *Stentors* by Vance Tartar and Gotram Uhlig. When grafting created cortical boundaries with a relatively weak stripe contrast (e.g. obtained by juxtaposition of the left and right cortex after removal of most of either the dorsal or ventral surface), oral development was delayed and preceded by a period of remodeling along the heal lines that involved branching of a subset of wider stripes that generated narrowest stripes and consequently created a stronger structural contrast ^43,46^ (reviewed in ^13^). Thus, it appears that left and right cortex interact to generate cues that remodel the cortex on both sides of the oral meridian. Double mutants lacking both Bcd1 and Hpo1, show defects not only in the positioning but also in the internal organization (including the left-right polarity of oral structures). Our observations suggest that Bcd1 and Hpo1 are parts of the left-right cross-talk that positions and shapes the ventral features including the oral apparatus.

To summarize, our observations suggest a multi-domain model for circumferential pattern formation in ciliates (Fig. 1C). Multiple cortical domains are formed by bilateral gradients of patterning factors including Hpo1, Bcd1 and Janus gene products. Adjacent or overlapping domains interact to regulate each other’s distribution and to create boundary effects that generate positional information for forming organelles.

## Materials and methods

### Strains and culture

All strains used were obtained from the *Tetrahymena* Stock Center (Cornell University, Ithaca NY, currently housed at Washington University, St Louis, MO USA; https://sites.wustl.edu/tetrahymena/). Cultures were grown in the SPPA medium (1% Proteose-peptone, 0.2% dextrose, 0.1% yeast extract and 0.003% EDTA:ferric sodium salt) supplemented with antibiotics (SPPA) ^50,51.^ CU428 *mpr1-1/mpr1-1* (*MPR1; VII*)(TSC_SD00178) and B2086 *mat1-2/mat1-2* (*mat1-2; II*) (TSC_SD00709) were used as wild type controls. CU427 *chx1-1/chx1-1* (*CHX1; VI*) (TSC_SD00715) was used for outcrosses to map the *hpo1* alleles. B*VII (TSC_SD00023) was used for self-crosses The following mutant strains were used: IA393 *hpo1-1/hpo1-1; eja1-1/eja1-1* (*hpo1-1, eja1-1; II*) (TSC_SD01463), IA418 *hpo1-2/hpo1-2* (*hpo1-2, VI*) (TSC_SD_01465), IA443 *hpo1-3/hpo1-3* (*hpo1-3; II*) (TSC_SD01455),, IA480 *hpo1-4/hpo1-4* (*hpo1-4, II*) (TSC_SD01466), IA359 *janC-1/janC-1 (janC-1, IV)* (TSC_SD00637), IA479 *janC-4/janC-4* (*janC-4; VII*) (TSC_SD01520), IA342 *bcd1-1/bcd1-1*, *eja1-1/eja1-1* (*bcd1-1, eja1-1*; II) (TSC_SD00635), IA378 *bcd1-2/bcd1-2* (*bcd1-2*, V) (TSC_SD00641), IA437 *janC-1/janC-1*, *hpo1-2/hpo1-2* (*janC-1, hpo1-2(3); IV*) (TSC_SD01576) and IA441 *bcd1-1/bcd1-1*, *hpo1-2/hpo1-2* (*bcd1-1, hpo1-2; IV*). In addition the following strains were made by editing the micronucleus by DNA homologous recombination (see below) and using crosses: UG20 *hpo1-F318S-3xHA-neo4/ hpo1-F318S-3xHA-neo4* (*hpo1-F318S-3xHA-neo4*), UG21 *hpo1::neo2/hpo1::neo2* (*hpo1::neo2*), and heterokaryon strains UG22 *bcd1-2/bcd1-2 hpo1::neo2/hpo1::neo2* (*BCD1, HPO1, pm-s*), UG23 *bcd1-2/bcd1-2 hpo1::neo2/hpo1::neo2* (*BCD1, HPO1, pm-s,* mates with UG22). To obtain progeny cells with the terminally lethal macronuclear genotype *hpo1::neo2*, *bcd1-2* in the macronucleus, UG22 and UG23 heterokaryon strains were starved and allowed to mate for 24 hr at 30°C, the cell population was incubated in SPPA medium for 6 hr followed by selection with paromomycin 100 μg/ml in SPPA to kill the parental cells.

### Mapping of hpo1 alleles and protein structure analysis

We applied ACCA method ^31^ to map the casual mutation for *hpo1-3* as described below (the same strategy was used to map the remaining three h*po1* alleles using appropriate homozygous strains). Strain IA443 was crossed to CU427 (homozygous for a cycloheximide (cy)-resistant allele *chx1-1* in micronucleus, TSC_SD00715). The heterozygous F1 progeny (*hpo1-3*/*HPO1*; *chx1-1/CHX1*) was assorted in SPPA to cycloheximide (cy) sensitivity. An assorted cy-sensitive F1 was mated to B*VII to produce F2 homozygotes using uniparental cytogamy ^52^. F2 clones were selected with 15 µg/ml cy and the cortical phenotype was evaluated by immunofluorescence. Twenty-three phenotypically wild-type or mutant F2 clones were pooled, cultured in 25ml of SPPA overnight, and subjected to starvation in 60 mM Tris-HCl (pH 7.5) for 2 days at 30°C. Total genomic DNA was extracted from the starved pools and used for generating genomic libraries using Illumina Truseq primer adapters. The libraries were sequence on Illumina HiSeq X to obtain paired-end 150 bp reads at 90x coverage. MiModD in the ACCA workflow was used to identify variants linked to the hpo1 phenotype as described in detail in ^29,31^. A 3D model of the Hpo1 structure was obtained from the AlphaFold protein structure Database (https://alphafold.com/). Protein domains were identified by InterPro (https://www.ebi.ac.uk/interpro/).

### Gene editing in T. thermophila

To construct a plasmid for a genomic knockout of *TTHERM_001276421/HPO1*, fragments were amplified from the genomic DNA of the wild-type strain CU428 and subcloned on the sides of the *neo4* selectable marker using primer pairs: 5’-AATTCCGCGGCGAACTTC TGAGTCATCATTG-3’, 5’-AATTCTGCAGCTTAAAGGCGTCTACCATTTTATTC-3’ and 5’-ATTCCCGGGTCAAGTATTCAACTCCTCTAAGTG-3’, 5’-AATTGGGCCCGAATACTCTGCTCGTGATGTCGA-3’. The resulting plasmid pcand2-KO-neo4 targeted for deletion a portion of the coding region composed of 1383 bp starting at codon 250 and including 371 bp of the 3’ UTR. Another targeting plasmid, pIA443-3HA was constructed to introduce the F318S substitution (*hpo1-3*) into the wild type background and simultaneously add a 3xHA epitope tag sequence at the 3’ end of the coding region. To this end, fragments were amplified using the total genomic DNA from strain IA443 (homozygous for *hpo1-3*) using primers pairs: 5’-AATTCTCGAGGAACTGTATTAGGAGTGATTTCAC-3’, 5’-TTAACTGCAGCATAAAATATCCAATTAATTTCAAATATCCACAAATATTATC-3’ and 5’-AATTGGATCCTTAACTAACTTCATCCTAGAAGCATTC-3’, 5’-AAATACGCGTTTAGTTCAACACTTAGAGGAGTAGAATAC-3’ and used to replace gene targeting parts of the plasmid pIFT54-3HA-native-neo4 ^53^. The targeting fragments of the above plasmids were released using restriction enzymes cleaving near the ends of flanking homologous sequences and the digested DNA was used for biolistic bombardment of mating CU428 and B2086 cells at an early stage of conjugation optimal for targeting in the micronucleus, followed by standard crosses to make mutant heterokaryons and homokaryons ^54^.

To prepare a strain for live imaging of Hpo1, the coding region of *TTHERM_001276421/HPO1* was cloned into the plasmid pGFP_PLK2-BTU1ov, downstream of the *MTT1* promoter and the GFP sequence. The primer pairs used to amplify the Hpo1 fragment were 5’-CTATACAAACGCGTGATGTAAAACTTACCTGATTGC-3’, and 5’-GTTCGCTTACGGATCCTCATTAACTAACTTCATCCTAGAAGC-3’. The transgene was placed between UTR sequences of *BTU1* for targeting to this locus by homologous DNA recombination. The plasmid was digested by BamHI and MluI, biolistically introduced into the CU428 strain and transformants were selected with 100 μg/ml paromomycin.

To overexpress Hpo1-HA, a plasmid was made (pMTT1_Hpo1_HA) with the following sequence elements cloned between the UTR sequences of the *BTU1* gene: *Bsr* selectable marker, *MTT1* promoter, coding region of HPO1 amplified with primers: 5’-TAAAATAATGGCCAAGTCGACAATGTAAAACTTACCTGATTGCG-3’ and 5’-AACATCATAAGGATAAGCACCGGATCCTTAACTAACTTCATCCTAGAAGCATTC-3’, HA epitope tag sequence. The targeting portion of the plasmid was released with SacI and BamHI, introduced biolistically into the *BTU1* locus (of CU428 strain) and transformants were selected with 60 μg/ml blastidicin S. To induce overproduction, the transgene-carrying cells were exposed to 2.5 μg/ml cadmium chloride for 6 hr.

### Microscopic imaging

*T. thermophila* cells were fixed by and prepared for immunofluorescence as described ^29,51^. The primary antibodies used were: anti-GFP (Rockland Immunochemicals, #600-401-215; 1:800 dilution), monoclonal anti-HA 16B12 (Covance; 1:300), and monoclonal anti-centrin 20H5 (EMD Millipore; 1:200-300;^55^). The secondary antibodies were conjugated to either Cy3 or FITC (Jackson ImmunoResearch, 115-095-146 and 111-165-003; 1:100–1:300). The nuclei were stained with DAPI (Sigma-Aldrich). The labeled cells were mounted in 90% glycerol, 10% PBS supplemented with 100 mg/ml DABCO (Sigma-Aldrich). To image the apical surface of cells, cell fragments were obtained as follows: 1.5 ml of cell culture was concentrated at 2,800 rpm/3 min, and washed with 1 ml of the nuclear isolation medium A (0.1M sucrose, 4% gum arabic, 0.0015M MgCl_2_, 0.01% spermidine-HCl, pH 6.75) ^56^. The cells were concentrated by centrifugation to 150 µl and combined with 160 μl of 1% paraformaldehyde/0.25% Triton X-100 followed by addition of 1.92 μl of octyl alcohol. The mixture was vortexed for 10-60 seconds and 20 μl of the sample was air-dried at the room temperature on a cover glass. Next, immunofluorescence was conducted as described above. The microscope images were collected on a Zeiss LSM 710 confocal microscope with a Plan-Apochromat 63×/1.40 oil DIC M27 objective and on an ELYRA S1 SR-SIM microscope equipped with a 63× NA 1.4 Oil Plan-Apochromat DIC. TIRF microscopy was executed as previously described ^57^ except that partial immobilization of cells was achieved by entrapment in a small volume of culture medium.

### Statistical analysis

Using the GraphPad PRISM software, we executed two-tailed unpaired t-tests to evaluate differences. P<0.05 was deemed statistically significant.

## Acknowledgements

This work was supported by the National Institutes of Health (grant R01GM135444 to J.G., R01GM110413 to K.F.L.), National Science Foundation (grant 1947608 to E.C.) and the German Federal Ministry of Education and Research (Bundesministerium für Bildung und Forschung) (grant 031L0101C de.NBI-epi to W.M.). The SR-SIM imaging was done at the Biomedical Microscopy Core, University of Georgia.

## Figure Legends

**Figure S1.**
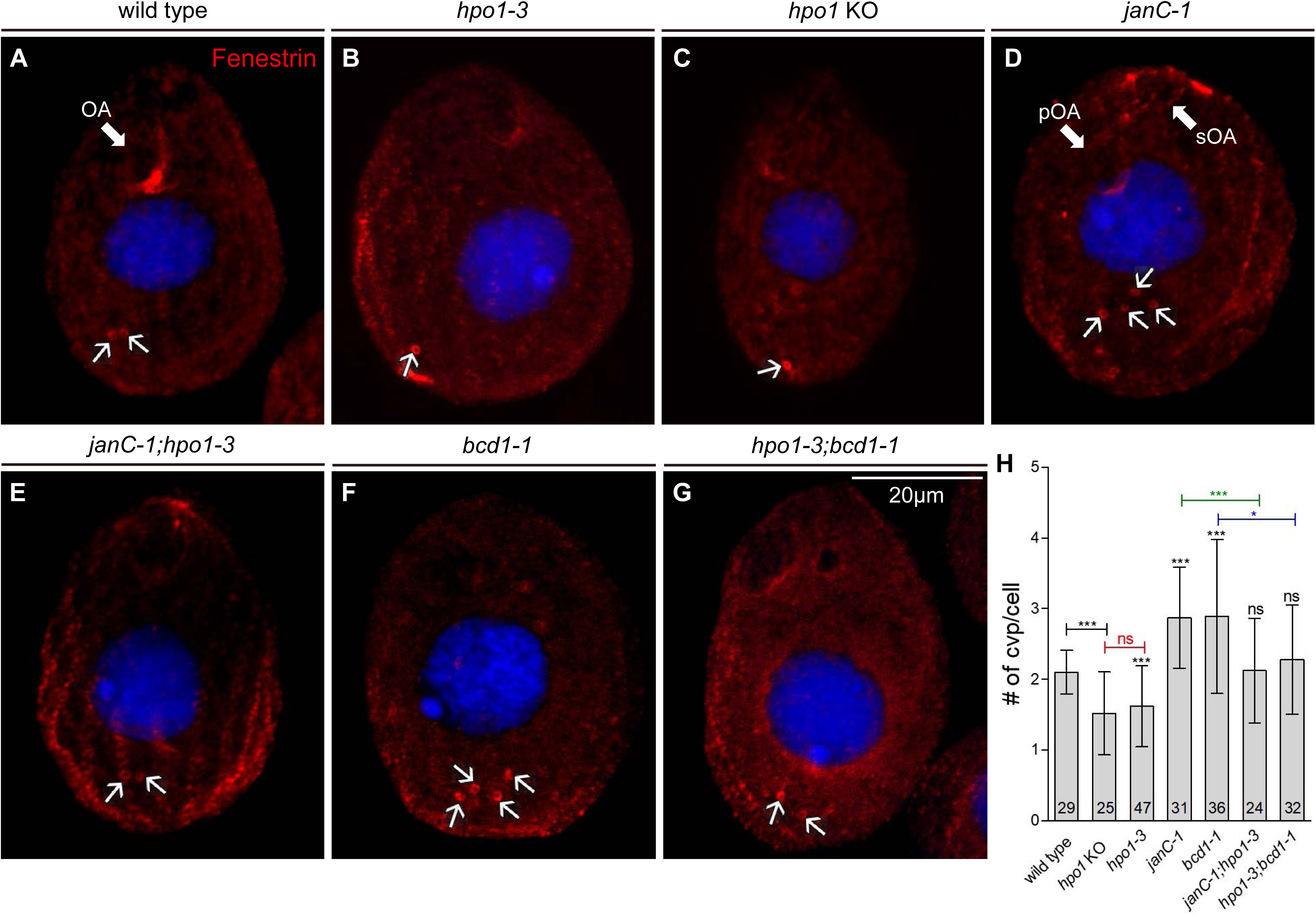
Quantification of the number of CVPs per cell. All genotypes are homozygous as indicated in the figure panels. (A-G) Confocal images of wild type (A), single mutants (B-D,F), and double mutants (E,G). The cells were labeled with the anti-fenestrin antibody (red), and DAPI (blue) after incubation for 4 hours at 38°C. Panels are representative of each genotype. The white arrows point to the CVPs. (H) The graph quantifies the number of CVPs per cell. The bars represent the means ± SD. The number of cells scored is displayed in each of the bar. A two-tailed unpaired t-test was executed for statistical analysis. ns: not significant. Stars indicate statistically significant (*: P < 0.05, **: P < 0.01, and ***: P < 0.001). Abbreviations: pOA, primary OA; sOA, secondary OA (in *janC-1* background).

**Figure S2.**
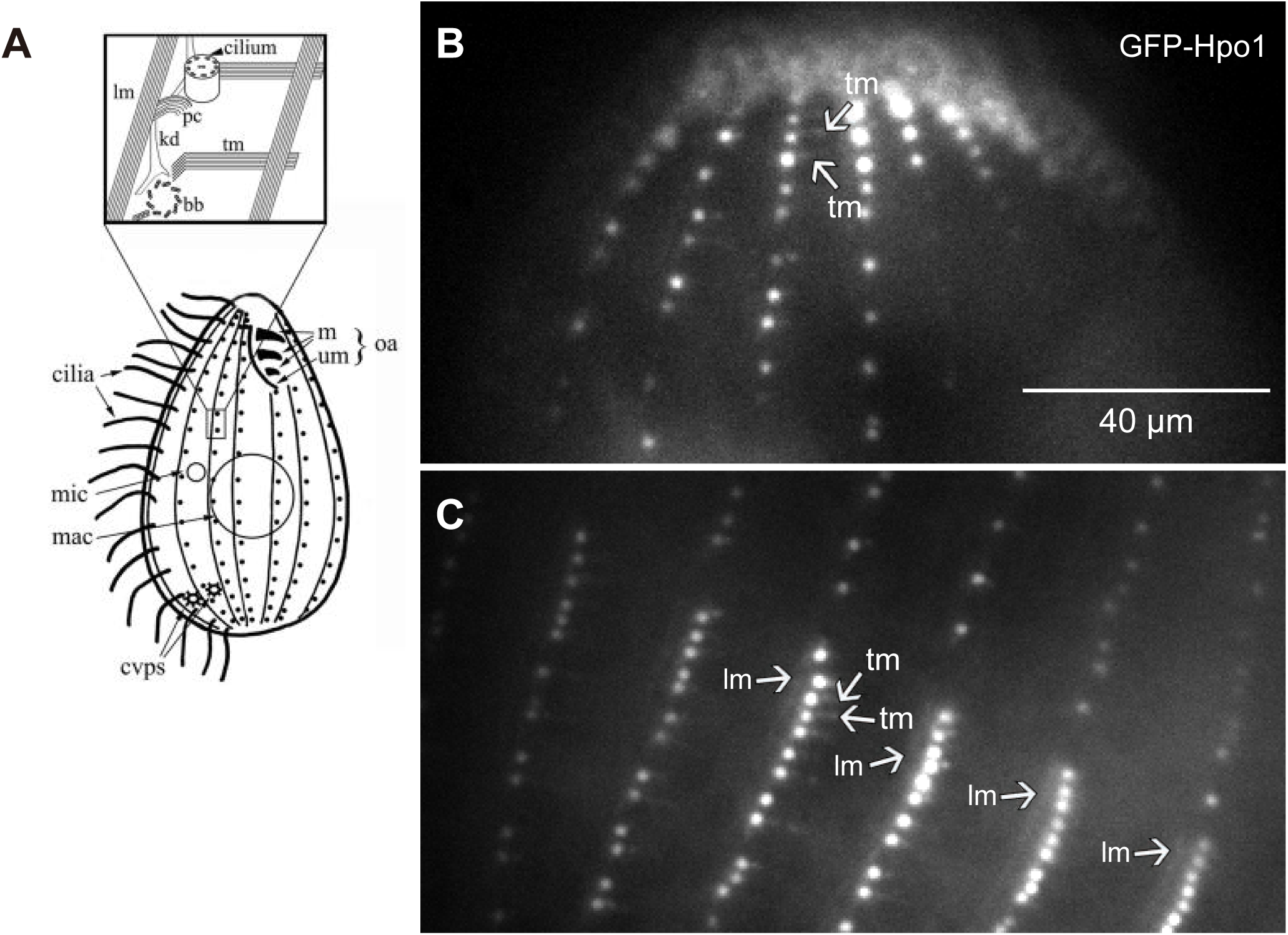
*In vivo* Hpo1 marks basal bodies and adjacent microtubule bundles. (A) Illustration of *T. thermophila* with the magnified inset describing a portion of cell cortex (from ^58^). (B,C) Still images of live cells expressing GFP-Hpo1 that localizes at positions consistent with the basal bodies and microtubule bundles (indicated by arrows). Abbreviations: tm, transverse microtubule; lm, longitudinal microtubule; pc, postciliary microtubule; bb, basal body, kd, kinetodesmal fiber (nonmicrotubular), m, membranelle; um, undulating membrane; oa, oral apparatus; mic, micronucleus; mac, macronucleus; cvp(s), contractile vacuole pore(s).

**Figure S3.**
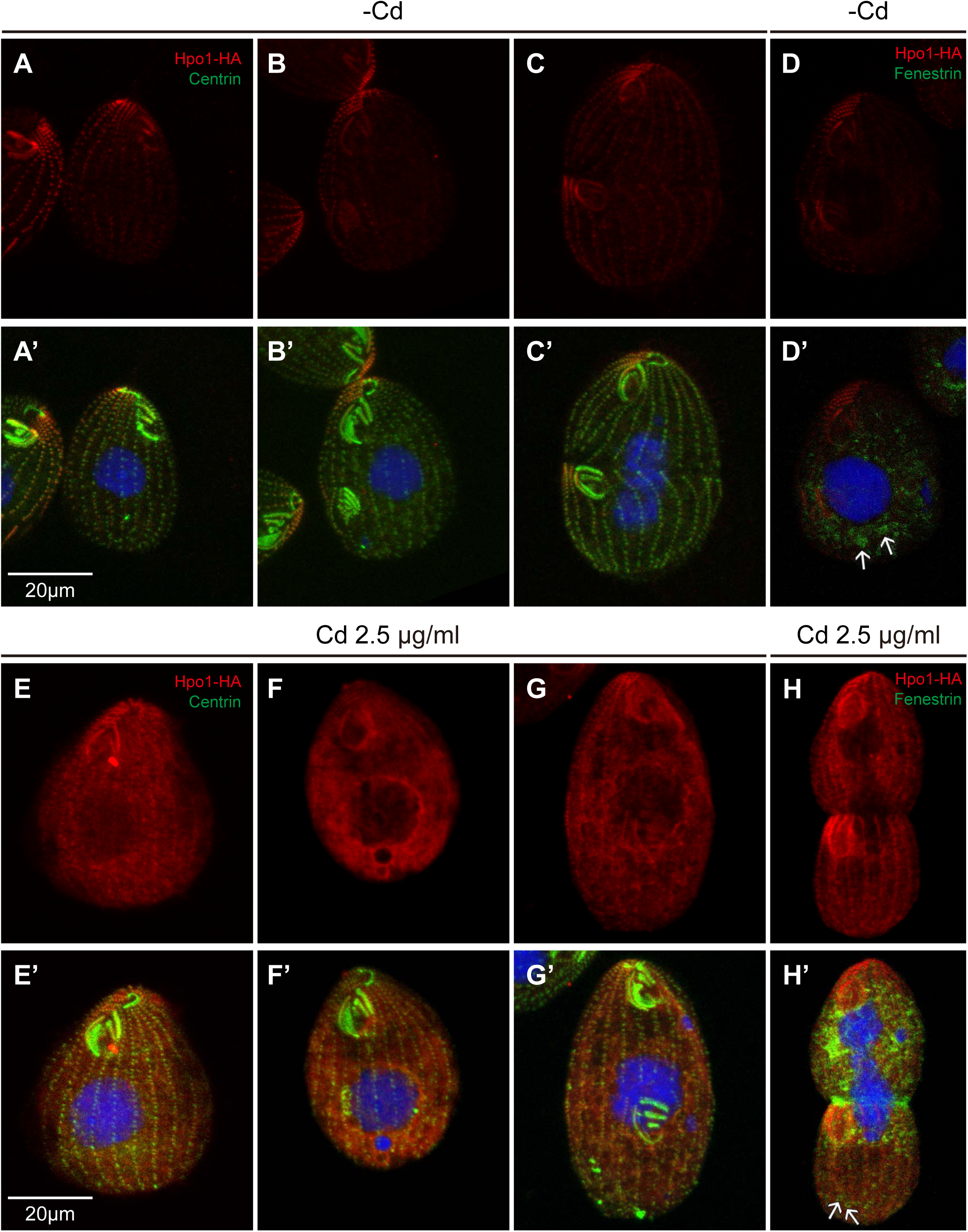
Overexpression of Hpo1 does not affect the cortical organelle pattern. Cells carrying a transgene expressing Hpo1-HA under the *MTT1* promoter were grown either without (A-D’) or with addition of 2.5 μg/ml cadmium chloride for 6 hr (E-H’), fixed and labeled with the anti-HA (red) and either anti-centrin (A’-C’; E’-G’) or anti-fenestrin (D’,H’) (green) antibodies and DAPI (blue). Note an accumulation of Hpo1-HA in the cell body of overproducing cells. Despite overproduction, the right-side and anterior gradients of Hpo1 are still apparent and the positions and number of OPs (E-G’ compare to A-C’) and the number of CVPs (H,H’ compare to D,D’) are unaffected.

**Figure S4.**
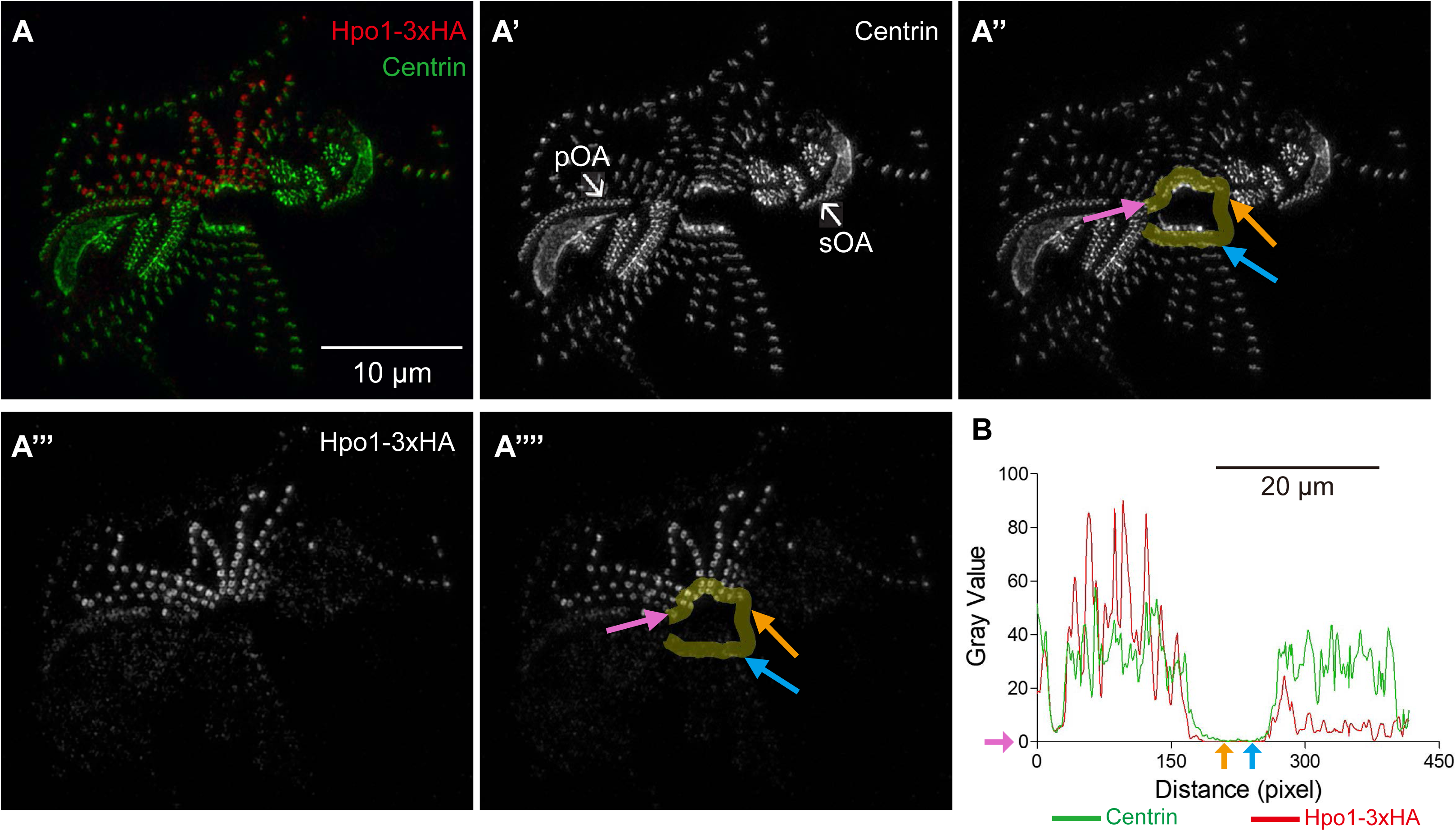
Quantification of the circumferential distribution of Hpo1-3xHA in the *janC-4* homozygote. (A-A’’’’) An SR-SIM image of Hpo1-3xHA and centrin in an apical cell fragment of the *janC-4* homozygote. (A) The cell fragment was labeled with anti-HA (red), 20H5 anti-centrin antibody (green), and DAPI (blue) after overnight incubation at 30°C. Duplicates of single channel grey scale images for centrin (A’,A’’) and Hpo1-3-HA (A’’’,A’’’’) are shown. The yellow lines mark the area used for measurements of signal intensity. The pink arrows show the start locations for signal intensity measurements. The orange and cyan arrow indicates initial and end point of gap, respectively. (B) The graph shows that signal intensity plots for Hpo1-3xHA (red line) and centrin (green line).

**Figure S5.**
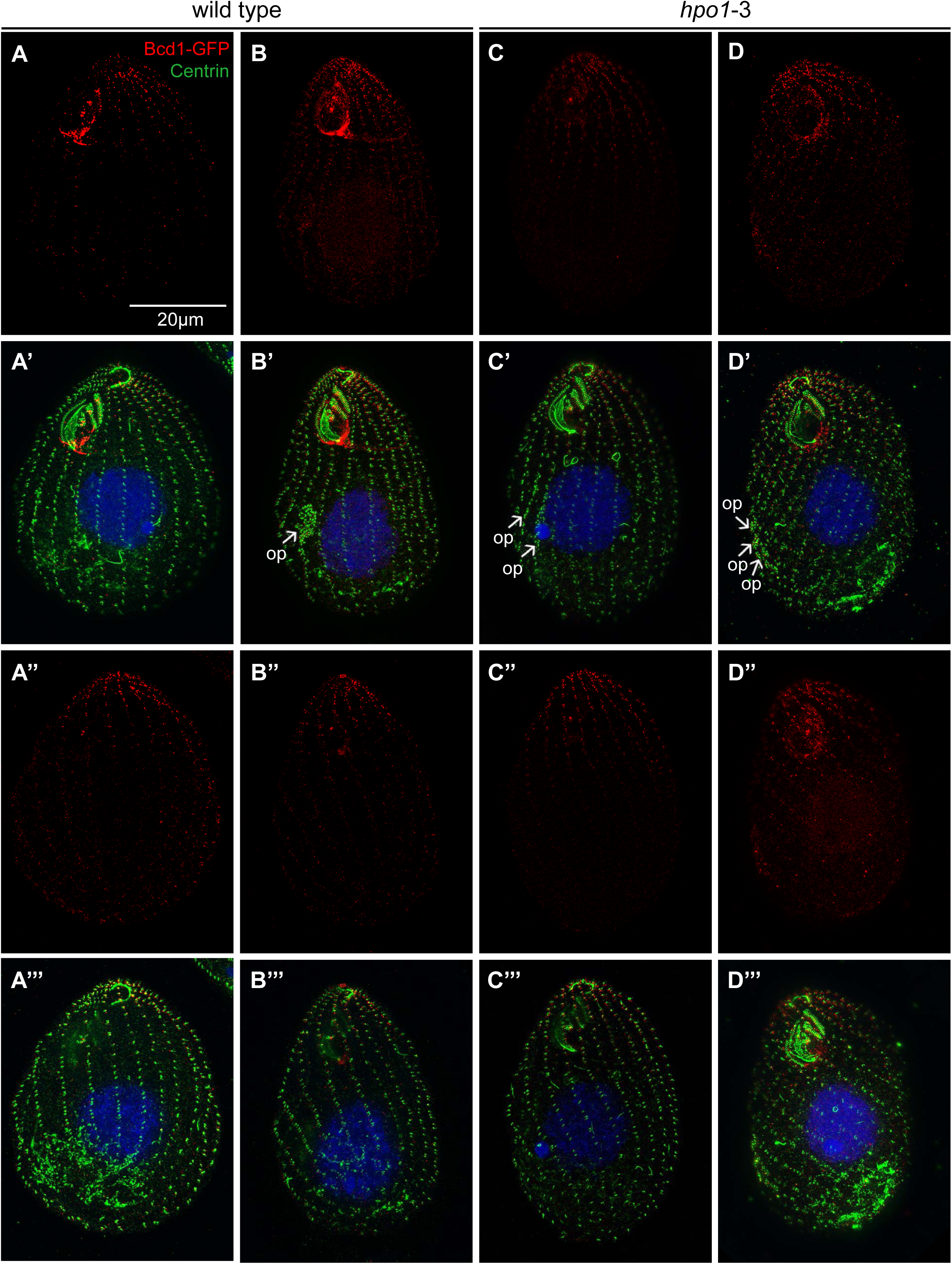
Localization of Bcd1-GFP in *hpo1-3 homozygotes*. (A-D’’’) Pairs of SR-SIM images showing two sides of the same cells that express Bcd1-GFP and are either otherwise wild-type (A-B’’’) or *hpo1-3* (C-D’’’). The cells were labeled with the anti-GFP antibodies (red), 20H5 anti-centrin antibody (green), and DAPI (blue) after a period of growth for 4 hours at 39°C.

**Figure S6.**
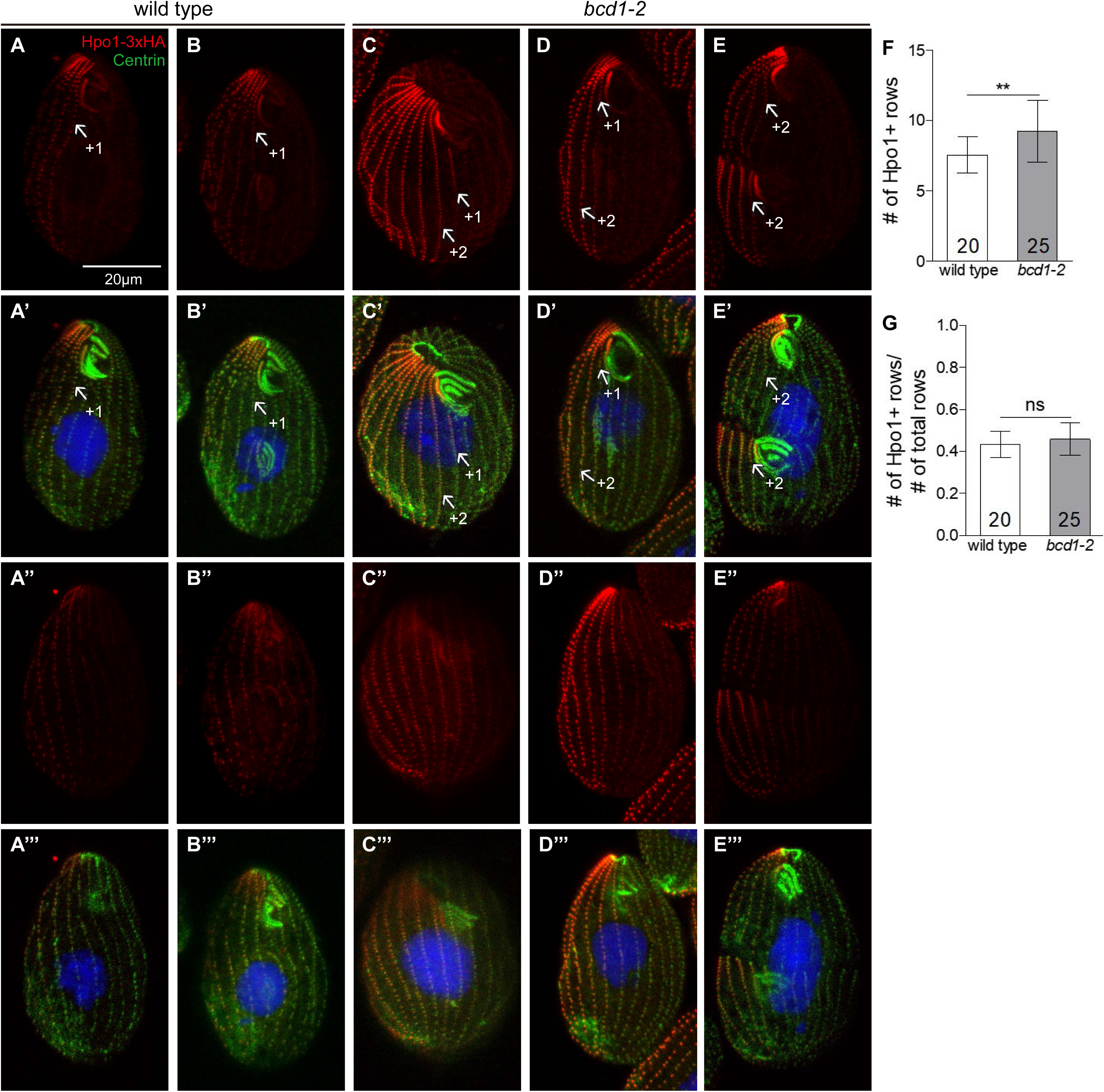
Comparison of Hpo1-3xHA distributions between the wild type and *bcd1-2* homozygotes. (A-E’’’) Confocal images of Hpo1-3xHA expressed in wild type (A-B’’’) or *bcd1-2* homozygotes (C-E’’’). The same images for *bcd1-2* background were also used for the Figure 8. After overnight incubation at 30°C, the cells were labeled with anti-HA antibodies (red), 20H5 anti-centrin antibody (green), and DAPI (blue). (F,G) Graphs quantify the number of rows enriched in Hpo1-3xHA (F), and the ratio of Hpo1 enriched rows to the total number of rows per cell (G). ns: not significant. Stars indicate statistically significant (*: P < 0.05, **: P < 0.01, and ***: P < 0.001).

**Figure S7.**
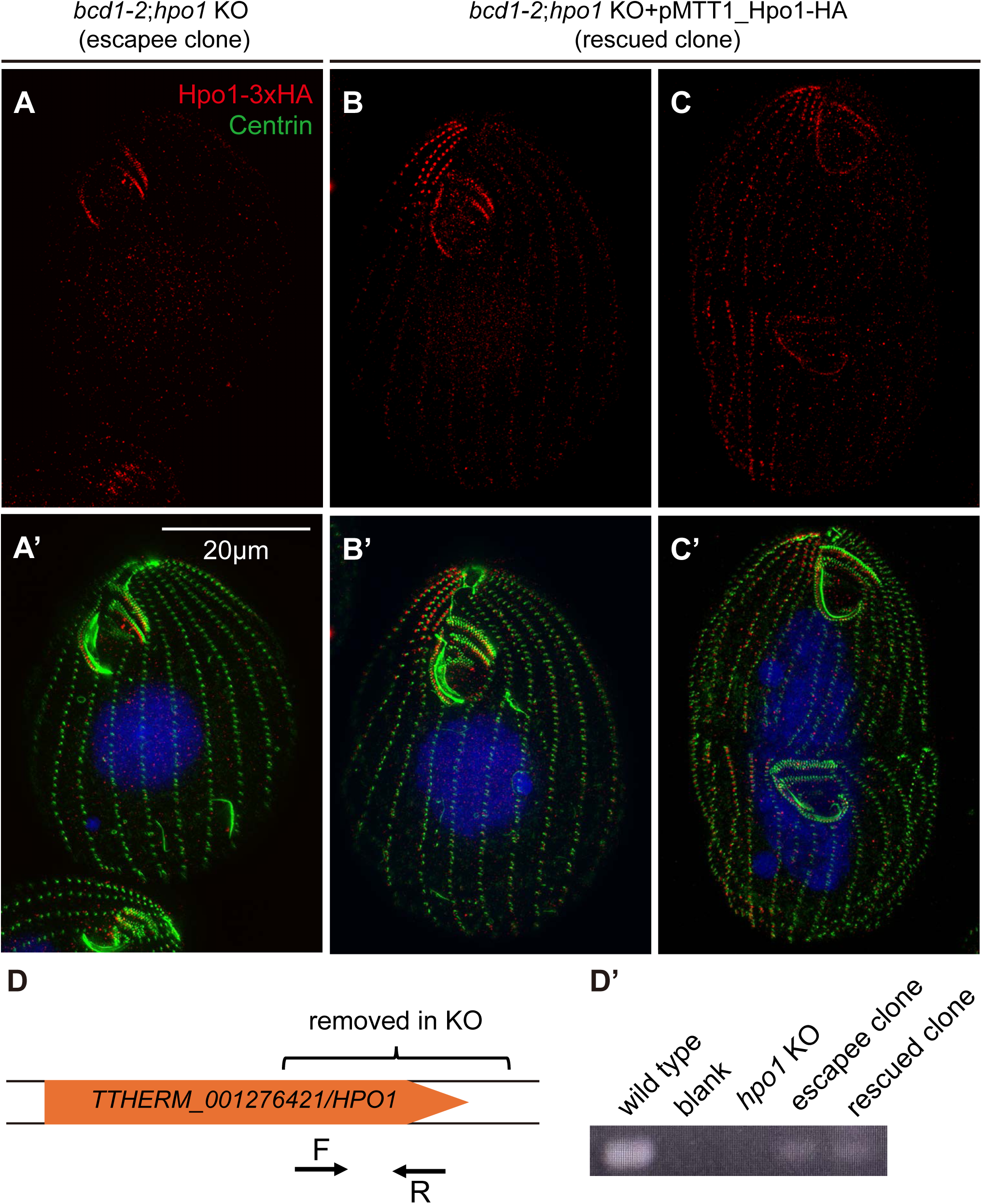
The lethality caused by the loss of function of both Bcd1 and Hpo1 can be rescued by a transgene encoding Hpo1-3xHA. (A-C’) SR-SIM images clones selected from the population of mating heterokaryons homozygous in the micronucleus for *bcd1-2* and *hpo1*-KO alleles that were either subjected to a mock biolistic bombardment (without transgene DNA) (A,A’) or biolistically transformed with a transgene encoding MTT1-Hpo1-3xHA (B-C’). The cells were labeled with anti-HA (red), 20H5 anti-centrin antibody (green), and DAPI (blue) after overnight incubation at 30°C. (D,D’) The diagram shows the positions of PCR primers designed for amplification of the portion of *HPO1* gene sequence deleted in the *hpo1-KO* allele (D) and the gel image showing the PCR products amplified from the genomic DNA isolated either from the escapee clone or from rescued clone (D’).

## References

1. Cole, E., and Gaertig, J. (2022). Anterior-posterior pattern formation in ciliates. J Eukaryot Microbiol, e12890. 10.1111/jeu.12890.

2. Ishida, T., Kaneko, Y., Iwano, M., and Hashimoto, T. (2007). Helical microtubule arrays in a collection of twisting tubulin mutants of Arabidopsis thaliana. Proc Natl Acad Sci U S A 104, 8544–8549. 10.1073/pnas.0701224104.

3. Hozumi, S., Maeda, R., Taniguchi, K., Kanai, M., Shirakabe, S., Sasamura, T., Speder, P., Noselli, S., Aigaki, T., Murakami, R., and Matsuno, K. (2006). An unconventional myosin in Drosophila reverses the default handedness in visceral organs. Nature 440, 798–802. 10.1038/nature04625.

4. Hatori, R., Ando, T., Sasamura, T., Nakazawa, N., Nakamura, M., Taniguchi, K., Hozumi, S., Kikuta, J., Ishii, M., and Matsuno, K. (2014). Left-right asymmetry is formed in individual cells by intrinsic cell chirality. Mech Dev 133, 146–162. 10.1016/j.mod.2014.04.002.

5. Speder, P., Adam, G., and Noselli, S. (2006). Type ID unconventional myosin controls left-right asymmetry in Drosophila. Nature 440, 803–807. 10.1038/nature04623.

6. Inaki, M., Liu, J., and Matsuno, K. (2016). Cell chirality: its origin and roles in left-right asymmetric development. Philos Trans R Soc Lond B Biol Sci 371. 10.1098/rstb.2015.0403.

7. Cole, E.S., Maier, W., Joachimiak, E., Jiang, Y.Y., Lee, C., Collet, E., Chmelik, C., Romero, D.P., Chalker, D., Alli, N.K., et al. (2023). The Tetrahymena bcd1 mutant implicates endosome trafficking in ciliate, cortical pattern formation. Mol Biol Cell 34, ar82. 10.1091/mbc.E22-11-0501.

8. Cole, E.S., Maier, W., Huynh, H.V., Reister, B., Sowunmi, D.O., Chukka, U.N., Lee, C., Parra, M., and Gaertig, J. (2024). Janus A : a polo kinase whose loss creates a dorsal/ventral intracellular homeosis in the ciliate, Tetrahymena.

9. Nelsen, E.M., and Frankel, J. (1989). Maintenance and regulation of cellular handedness in Tetrahymena. Development 105, 457–471. 10.1242/dev.105.3.457.

10. Nelsen, E.M., Frankel, J., and Williams, N.E. (1989). Oral assembly in left-handed Tetrahymena thermophila. J Protozool 36, 582–596. 10.1111/j.1550-7408.1989.tb01101.x.

11. Yao, M.C., and Yao, C.H. (1991). Transformation of Tetrahymena to cycloheximide resistance with a ribosomal protein gene through sequence replacement. Proc Natl Acad Sci U S A 88, 9493–9497. 10.1073/pnas.88.21.9493.

12. Galati, D.F., Bonney, S., Kronenberg, Z., Clarissa, C., Yandell, M., Elde, N.C., Jerka-Dziadosz, M., Giddings, T.H., Frankel, J., and Pearson, C.G. (2014). DisAp-dependent striated fiber elongation is required to organize ciliary arrays. J Cell Biol 207, 705–715. 10.1083/jcb.201409123.

13. Frankel, J. (1989). Pattern formation : ciliate studies and models (Oxford University Press).

14. Frankel, J. (2008). What do genic mutations tell us about the structural patterning of a complex single-celled organism? Eukaryot Cell 7, 1617–1639. 10.1128/EC.00161-08.

15. Cole, E.S., Frankel, J., and Jenkins, L.M. (1987). bcd: A mutation affecting the width of organelle domains in the cortex of Tetrahymena thermophila. Rouxs Arch Dev Biol 196, 421–433. 10.1007/BF00399142.

16. Frankel, J., Jenkins, L.M., Nelsen, E.M., and Stoltzman, C.A. (1993). Hypoangular: a gene potentially involved in specifying positional information in a ciliate, Tetrahymena thermophila. Dev Biol 160, 333–354. 10.1006/dbio.1993.1311.

17. Louka, P., Vasudevan, K.K., Guha, M., Joachimiak, E., Wloga, D., Tomasi, R.F., Baroud, C.N., Dupuis-Williams, P., Galati, D.F., Pearson, C.G., et al. (2018). Proteins that control the geometry of microtubules at the ends of cilia. J Cell Biol 217, 4298–4313. 10.1083/jcb.201804141.

18. Van De Weghe, J.C., Rusterholz, T.D.S., Latour, B., Grout, M.E., Aldinger, K.A., Shaheen, R., Dempsey, J.C., Maddirevula, S., Cheng, Y.H., Phelps, I.G., et al. (2017). Mutations in ARMC9, which Encodes a Basal Body Protein, Cause Joubert Syndrome in Humans and Ciliopathy Phenotypes in Zebrafish. Am J Hum Genet 101, 23–36. 10.1016/j.ajhg.2017.05.010.

19. Latour, B.L., Van De Weghe, J.C., Rusterholz, T.D., Letteboer, S.J., Gomez, A., Shaheen, R., Gesemann, M., Karamzade, A., Asadollahi, M., Barroso-Gil, M., et al. (2020). Dysfunction of the ciliary ARMC9/TOGARAM1 protein module causes Joubert syndrome. J Clin Invest 130, 4423–4439. 10.1172/JCI131656.

20. Allen, R.D. (1969). The morphogenesis of basal bodies and accessory structures of the cortex of the ciliated protozoan Tetrahymena pyriformis. J Cell Biol 40, 716–733. 10.1083/jcb.40.3.716.

21. Nelsen, E.M., Williams, N.E., Yi, H., Knaak, J., and Frankel, J. (1994). "Fenestrin" and conjugation in Tetrahymena thermophila. J Eukaryot Microbiol 41, 483–495. 10.1111/j.1550-7408.1994.tb06047.x.

22. Cole, E.S., Anderson, P.C., Fulton, R.B., Majerus, M.E., Rooney, M.G., Savage, J.M., Chalker, D., Honts, J., Welch, M.E., Wentland, A.L., et al. (2008). A proteomics approach to cloning fenestrin from the nuclear exchange junction of tetrahymena. J Eukaryot Microbiol 55, 245–256. 10.1111/j.1550-7408.2008.00337.x.

23. Shang, Y., Song, X., Bowen, J., Corstanje, R., Gao, Y., Gaertig, J., and Gorovsky, M.A. (2002). A robust inducible-repressible promoter greatly facilitates gene knockouts, conditional expression, and overexpression of homologous and heterologous genes in Tetrahymena thermophila. Proc Natl Acad Sci U S A 99, 3734–3739. 10.1073/pnas.052016199.

24. Frankel, J., and Nelsen, E.M. (1986). How the mirror-image pattern specified by a janus mutation of Tetrahymena thermophila comes to expression. Dev Genet 6, 213–238. 10.1002/dvg.1020060306.

25. Frankel, J., and Nelsen, E.M. (1987). Positional reorganization in compound janus cells of Tetrahymena thermophila. Development 99, 51–68. 10.1242/dev.99.1.51.

26. Frankel, J., and Jenkins, L.M. (1979). A mutant of Tetrahymena thermophila with a partial mirror-image duplication of cell surface pattern. II. Nature of genic control. J Embryol Exp Morphol 49, 203–227.

27. Jerka-Dziadosz, M., and Frankel, J. (1979). A mutant of Tetrahymena thermophila with a partial mirror-image duplication of cell surface pattern. I. Analysis of the phenotype. J Embryol Exp Morphol 49, 167–202.

28. Orias, E., and Rasmussen, L. (1976). Dual capacity for nutrient uptake in Tetrahymena. IV. Growth without food vacuoles and its implications. Exp Cell Res 102, 127–137. 10.1016/0014-4827(76)90307-4.

29. Jiang, Y.Y., Maier, W., Chukka, U.N., Choromanski, M., Lee, C., Joachimiak, E., Wloga, D., Yeung, W., Kannan, N., Frankel, J., and Gaertig, J. (2020). Mutual antagonism between Hippo signaling and cyclin E drives intracellular pattern formation. J Cell Biol 219. 10.1083/jcb.202002077.

30. Tavares, A., Goncalves, J., Florindo, C., Tavares, A.A., and Soares, H. (2012). Mob1: defining cell polarity for proper cell division. J Cell Sci 125, 516–527. 10.1242/jcs.096610.

31. Jiang, Y.Y., Maier, W., Baumeister, R., Minevich, G., Joachimiak, E., Ruan, Z., Kannan, N., Clarke, D., Frankel, J., and Gaertig, J. (2017). The Hippo Pathway Maintains the Equatorial Division Plane in the Ciliate Tetrahymena. Genetics 206, 873–888. 10.1534/genetics.117.200766.

32. Jiang, Y.Y., Maier, W., Baumeister, R., Joachimiak, E., Ruan, Z., Kannan, N., Clarke, D., Louka, P., Guha, M., Frankel, J., and Gaertig, J. (2019). Two Antagonistic Hippo Signaling Circuits Set the Division Plane at the Medial Position in the Ciliate Tetrahymena. Genetics 211, 651–663. 10.1534/genetics.118.301889.

33. Slabodnick, M.M., Ruby, J.G., Dunn, J.G., Feldman, J.L., DeRisi, J.L., and Marshall, W.F. (2014). The kinase regulator mob1 acts as a patterning protein for stentor morphogenesis. PLoS Biol 12, e1001861. 10.1371/journal.pbio.1001861.

34. Lee, C., Maier, W., Jiang, Y.Y., Nakano, K., Lechtreck, K.F., and Gaertig, J. (2024). Global and local functions of the Fused kinase ortholog CdaH in intracellular patterning in Tetrahymena. J Cell Sci 137. 10.1242/jcs.261256.

35. Breslow, D.K., Hoogendoorn, S., Kopp, A.R., Morgens, D.W., Vu, B.K., Kennedy, M.C., Han, K., Li, A., Hess, G.T., Bassik, M.C., et al. (2018). A CRISPR-based screen for Hedgehog signaling provides insights into ciliary function and ciliopathies. Nat Genet 50, 460–471. 10.1038/s41588-018-0054-7.

36. Cavalier-Smith, T., and Chao, E.E. (2004). Protalveolate phylogeny and systematics and the origins of Sporozoa and dinoflagellates (phylum Myzozoa nom. nov.). Eur J Protistol 40, 185–212. 10.1016/j.ejop.2004.01.002.

37. Janouskovec, J., Tikhonenkov, D.V., Mikhailov, K.V., Simdyanov, T.G., Aleoshin, V.V., Mylnikov, A.P., and Keeling, P.J. (2013). Colponemids Represent Multiple Ancient Alveolate Lineages. Current Biology 23, 2546–2552. 10.1016/j.cub.2013.10.062.

38. Orias, E. (1976). Derivation of Ciliate Architecture from a Simple Flagellate - Evolutionary Model. T Am Microsc Soc 95, 415–429. Doi 10.2307/3225135.

39. Tartar, V. (1961). The biology of Stentor (Pergammon Press).

40. Gruber, A. (1885). Ueber kiinstliche Teilung bei Infusorien. Biologisches Zentralblatt 3, 580–582.

41. Morgan, T.H. (1901). Regeneration of proportionate structures in stentor.

42. Tartar, V. (1956). IV. PATTERN AND SUBSTANCE IN STENTOR. In Cellular Mechanics in Differentiation and Growth, (Princeton University Press), pp. 73–100. doi:10.1515/9781400876877-005.

43. Uhlig, G. (1960). Entwicklungsphysiologische Untersuchungen zur Morphogenese von Stentor coeruleus Ehrbg. Arch. Protistenk. 105, 1–109.

44. Paulin, J.J., and Bussey, J. (1971). Oral regeneration in the ciliate Stentor coeruleus: a scanning and transmission electron optical study. J Protozool 18, 201–213. 10.1111/j.1550-7408.1971.tb03308.x.

45. Pelvat, B., and Dehaller, G. (1979). La Regeneration de L’appareil Oral chez Stentor coeruleus: Etude au Protargol et Essai de Morphogenese Comparee. Protistologica 15, 369–386.

46. Tartar, V. (1956). Grafting Experiments Concerning Primordium Formation in Stentor Coeruleus. Journal of Experimental Zoology 131, 75–121. DOI 10.1002/jez.1401310105.

47. Cole, E.S., Frankel, J., and Jenkins, L.M. (1988). Interactions between janus and bcd cortical pattern mutants in Tetrahymena thermophila : An investigation of intracellular patterning mechanisms using double-mutant analysis. Rouxs Arch Dev Biol 197, 476–489. 10.1007/BF00385681.

48. Frankel, J., Jenkins, L.M., and Bakowska, J. (1984). Selective mirror-image reversal of ciliary patterns in Tetrahymena thermophila homozygous for ajanus mutation. Wilehm Roux Arch Dev Biol 194, 107–120. 10.1007/BF00848350.

49. Frankel, J., Nelsen, E., and Jenkins, L. (1987). Intracellular pattern reversal in Tetrahymena thermophila: janus mutants and their geometrical phenocopies. pp. 219–244.

50. Gorovsky, M.A. (1973). Macro- and micronuclei of Tetrahymena pyriformis: a model system for studying the structure and function of eukaryotic nuclei. J Protozool 20, 19–25. 10.1111/j.1550-7408.1973.tb05995.x.

51. Gaertig, J., Wloga, D., Vasudevan, K.K., Guha, M., and Dentler, W. (2013). Discovery and functional evaluation of ciliary proteins in Tetrahymena thermophila. Methods Enzymol 525, 265–284. 10.1016/B978-0-12-397944-5.00013-4.

52. Cole, E.S., and Bruns, P.J. (1992). Uniparental cytogamy: a novel method for bringing micronuclear mutations of Tetrahymena into homozygous macronuclear expression with precocious sexual maturity. Genetics 132, 1017–1031. 10.1093/genetics/132.4.1017.

53. Hazime, K.S., Zhou, Z., Joachimiak, E., Bulgakova, N.A., Wloga, D., and Malicki, J.J. (2021). STORM imaging reveals the spatial arrangement of transition zone components and IFT particles at the ciliary base in Tetrahymena. Sci Rep 11, 7899. 10.1038/s41598-021-86909-5.

54. Dave, D., Wloga, D., and Gaertig, J. (2009). Manipulating ciliary protein-encoding genes in Tetrahymena thermophila. Methods Cell Biol 93, 1–20. 10.1016/S0091-679X(08)93001-6.

55. Salisbury, J.L., Baron, A.T., and Sanders, M.A. (1988). The centrin-based cytoskeleton of Chlamydomonas reinhardtii: distribution in interphase and mitotic cells. J Cell Biol 107, 635–641. 10.1083/jcb.107.2.635.

56. Gorovsky, M.A. (1970). Studies on nuclear structure and function in Tetrahymena pyriformis. 3. Comparison of the histones of macro- and micronuclei by quantitative polyacrylamide gel electrophoresis. J Cell Biol 47, 631-636. 10.1083/jcb.47.3.631.

57. Jiang, Y.Y., Lechtreck, K., and Gaertig, J. (2015). Total internal reflection fluorescence microscopy of intraflagellar transport in Tetrahymena thermophila. Methods Cell Biol 127, 445–456. 10.1016/bs.mcb.2015.01.001.

58. Thazhath, R., Jerka-Dziadosz, M., Duan, J., Wloga, D., Gorovsky, M.A., Frankel, J., and Gaertig, J. (2004). Cell context-specific effects of the beta-tubulin glycylation domain on assembly and size of microtubular organelles. Mol Biol Cell 15, 4136–4147. 10.1091/mbc.e04-03-0247.

